# Pulvinar perturbation causes functional reorganization of visual attentional information in posterior parietal cortex

**DOI:** 10.1101/2024.09.29.615686

**Authors:** Fang-Chi Yang, Feng-Kuei Chiang, Erin L. Rich, Rebecca D. Burwell

**Affiliations:** Cognitive and Psychological Sciences, Brown University, Providence, RI, 02912, USA; Nash Family Department of Neuroscience, Lipschultz Center for Cognitive Neuroscience and Friedman Brain Institute, Icahn School of Medicine at Mount Sinai, New York, NY 10029, USA

**Keywords:** lateral posterior nucleus, neural ensemble, population coding, decoding, spatial tuning

## Abstract

The pulvinar nucleus of the thalamus (PUL) is a higher-order thalamic relay and the main visual extrageniculate thalamic nucleus. Evidence suggests the PUL coordinates information processing across cortical areas involved in visual perception and attention. Other findings suggest the PUL may also influence higher-order cognitive processes, such as attentional control, through interactions with connected neocortical areas like the posterior parietal cortex (PPC). We hypothesized that PUL input to the dorsal and caudal PPC (DPPC and CPPC) enhances visuospatial processing and attention. To test this hypothesis, we recorded neuronal activity in the PUL, DPPC, and CPPC of freely behaving rats performing the visuospatial attention (VSA) task while optogenetically suppressing PUL neurons on some trials. We found that PUL manipulation did not affect behavioral performance, but reorganized neural codes in DPPC and CPPC as well as PUL itself.

## Introduction

The pulvinar nucleus of the thalamus (PUL), also called the lateral posterior nucleus in rodents, is a higher-order thalamic relay and the main visual extrageniculate thalamic nucleus. The PUL shows functional and connectional homology with the primate pulvinar. Both the rodent and primate PUL receive inputs from the superior colliculus (SC) and interconnect heavily with multiple cortical areas including visual cortex and higher order association regions^1–7^. Available evidence is consistent with the idea that the PUL serves as a modulator for coordinating information processing across multiple cortical areas involved in visual perception and attention^8–10^. New evidence, however, suggests the PUL may also play a role in higher order cognitive processes such as the control of attention through interactions with its connected neocortical areas^11^ ^12^. One such target of the PUL is the posterior parietal cortex (PPC). Both the PUL and the PPC have long been implicated in visual attention and visuospatial functions ^6,13–17^. However, how attention is modulated by PUL projections to the PPC is understudied.

Most previous studies of the rodent PPC focused on its dorsal portion, largely ignoring the ventral and caudal aspects of the region ^18–21^ (but see ^22,23^). We recently examined functional neuroanatomical and electrophysiological differences between dorsal PPC (DPPC) and caudal PPC (CPPC) using our novel visuospatial attention (VSA) task (Figure 1A-B) ^4^. In the VSA task, rats must employ controlled attention to monitor multiple locations for the onset of a target stimulus. The brief presentation of a target stimulus recruits stimulus-driven attention. Rats are then required to indicate the target location, and correct decisions are followed by a food reward. The VSA task engages perception, controlled and stimulus-driven attention, decision-making, and reward learning. The task also allows dissociation of neuronal activity correlated with egocentric and allocentric frames of reference. We previously showed that DPPC and CPPC neurons code for both egocentric and allocentric frames of reference. Importantly, we also provided evidence that the DPPC is more engaged in controlled attention and CPPC is more engaged in stimulus driven attention, a finding consistent with the functional differences in dorsal and ventral PPC in the primate brain ^4,24,25^.

**Figure 1.**
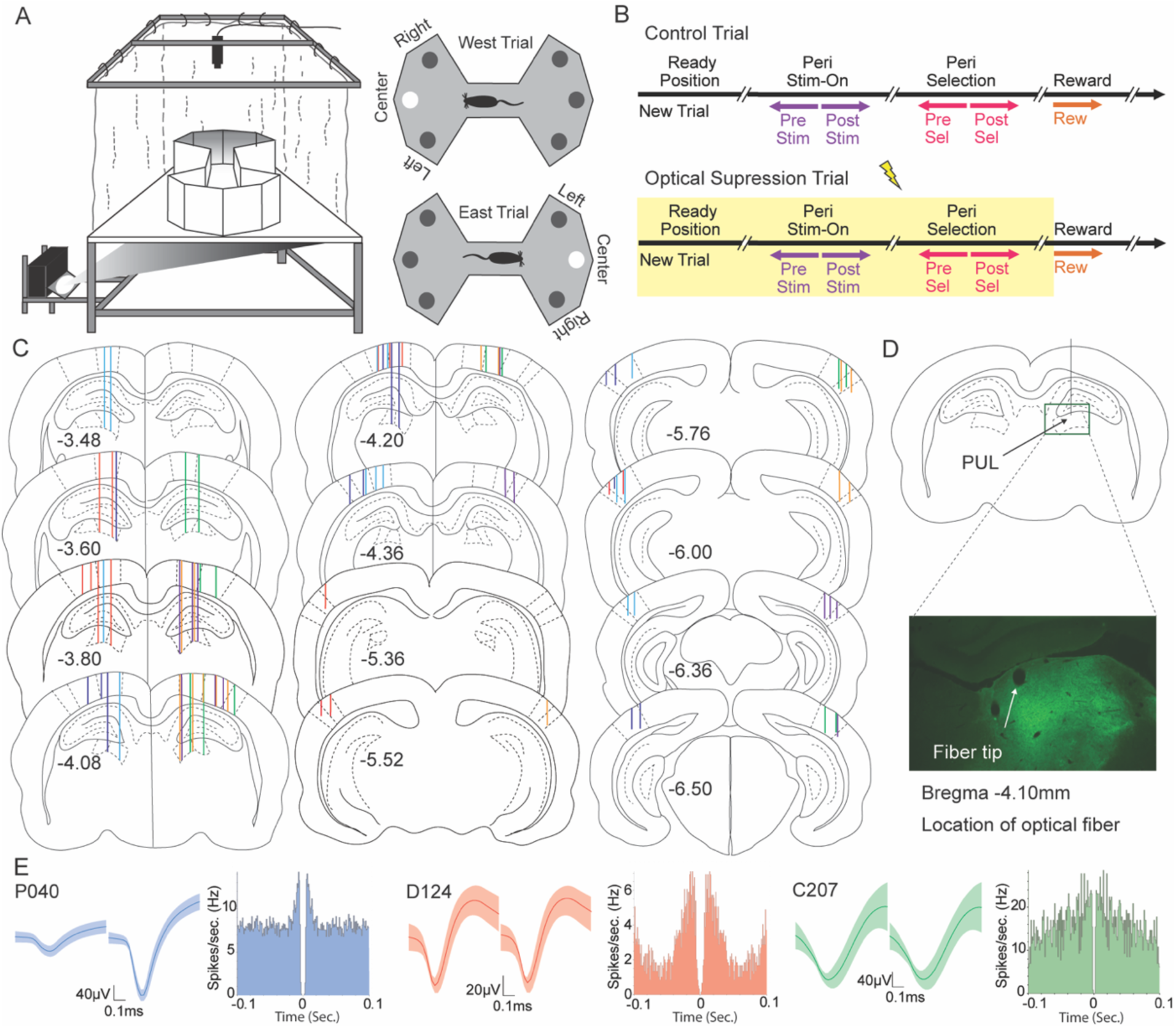
Multisite recording and optical perturbation in the Visuospatial Attention (VSA) task in the pulvinar (PUL), dorsal posterior parietal cortex (DPPC), and caudal posterior parietal cortex (CPPC). (A) Schematic of the Floor Projection Maze with a bowtie-shaped enclosure used for the VSA task (left). Top-down view of back-projected for west vs. east trials (right). Trials were initiated when the rat stopped in the middle of the maze facing west or east. After a variable period, one of three possible locations was briefly illuminated. The animal made a selection by approaching one of the three locations, and then returned to the center of the maze for a food reward. A new trial was immediately triggered on the alternate side. (B) Progression of a trial without or with optical suppression. Five epochs were analyzed for behavioral correlates, including the 500 ms prior to stimulus onset, after stimulus onset, prior to selection, after selection, and after the animal entered the reward zone. Optical suppression was randomly delivered to 50% of trials. Light delivery began when the rat entered the ready position initiating a new trial and ended when the rat entered the reward zone terminating the trial. (C) Estimated locations of implanted stereotrodes in either the left or right hemispheres. PUL and DPPC locations are shown from −3.48 to −4.36 mm relative to bregma, and CPPC locations are shown from −5.36 to −6.56 mm relative to bregma. Locations are color coded by subject: 14050, green; 14059, dark blue; 14060, red; 14061, marigold yellow; 14062, purple; 14064, sky blue. (D) Representative example of transduction of halorhodopsin in the pulvinar (PUL). (E) Example waveforms (mean ± stdev) and autocorrelograms (bin width1ms) for isolated cells in the PUL (left); DPPC (middle); and CPPC (right).

The DPPC and CPPC, which are considered higher-order visual associational regions, are robustly and reciprocally connected with the PUL. To probe the function of this circuitry, we recorded neuronal activity in the rat PUL during performance on the VSA task ^12^. PUL correlates of behavior in this task were similar to what we had observed in the DPPC and CPPC ^4,12,20^. PUL neurons signaled location in both egocentric and allocentric spatial frameworks. PUL neurons also correlated with selection behavior and signaled stimulus onset suggesting that the PUL is engaged visuospatial attention. One possibility is that the rodent PUL modulates both controlled attention and stimulus-driven through its reciprocal connections with dorsal and caudal PPC respectively ^4,6,7^. By this view, the PUL relays visual information to both DPPC and CPPC, which also receive visual information from other striate and extrastriate sources. The PUL input serves to amplify and coordinate visuospatial information in the dorsal and caudal PPC, which then may be projected back to the PUL through reciprocal connections for further processing and integration.

In this study, we combined multi-site recording and optogenetic methods to examine the role of the PUL connections with the DPPC and VPPC in visuospatial attention. We hypothesized that PUL input to the DPPC and CPPC enhances visuospatial information processing and improves visuospatial attention, such that PUL suppression should result in degraded representations of task-relevant information. PUL input may support the stimulus-driven functions of the CPPC and the controlled attention functions of the DPPC. If this is the case, then PUL suppression may alter neuronal correlates in the CPPC and the DPPC. To address these hypotheses, we simultaneously recorded neuronal activity in the PUL, DPPC, and CPPC of freely behaving rats performing the VSA task while optogenetically suppressing PUL neurons on some trials. We found that manipulation of the PUL modestly impacted behavioral performance, and only slightly reduced task-related neuronal correlates in both DPPC and CPPC with some indications that the CPPC was more affected. However, the impact was not always in the loss of selectivity. Although some cells did lose behavioral correlates with optical suppression, new selective cells emerged, and some cells altered patterns of selectivity, such that substantial coding for task relevant information was still evident in all three regions. Therefore, optical perturbation of the PUL resulted in the reorganization of information represented in the PUL itself as well as the downstream PPC regions.

## Results

Using a circuit analysis approach, we examined how the PUL interacts with the DPPC and CPPC under visuospatial attentional demands. We trained rats on our VSA task, which was adapted from the five-choice serial reaction time task ^26^ for use in our floor projection maze apparatus ^27,28^ (Figure 1A). Rats were required to approach and wait in the ready position in the center of the bowtie maze. After a variable time, a stimulus appeared for 500 msec in one of three egocentric locations (left, center, or right) on one side of the maze (east or west, Figure 1B). The rat made a selection by approaching the cued location, and a food reward was delivered in the center of the maze following a correct response. Trials alternated between the east and west sides of the maze. Failure to approach during the variable response window was considered an omission. Thus, trials were either correct, incorrect, or omitted. The chance level of performance on completed trials is 33.33%. The mean accuracy across 100 recording sessions from 6 rats was 64.3% (range 41.9%-90.6%).

### Neurons signaled stimulus, selection, outcome and location during control trials

We recorded a total of 474 PUL cells, 318 DPPC cells, and 225 CPPC cells from six rats that met single unit quality criteria for inclusion (Figure 1C-E). To understand basic response properties in this task, we first analyzed LED-OFF trials only (Figure 2). We first established neuronal correlates of each cell by comparing firing rates across conditions or epochs. We then analyzed neuronal correlates using a multiway chi-square test of independence. Because we were interested in regional differences, planned or post hoc pairwise comparisons were conducted using chi-square tests with Bonferroni correction.

**Figure 2.**
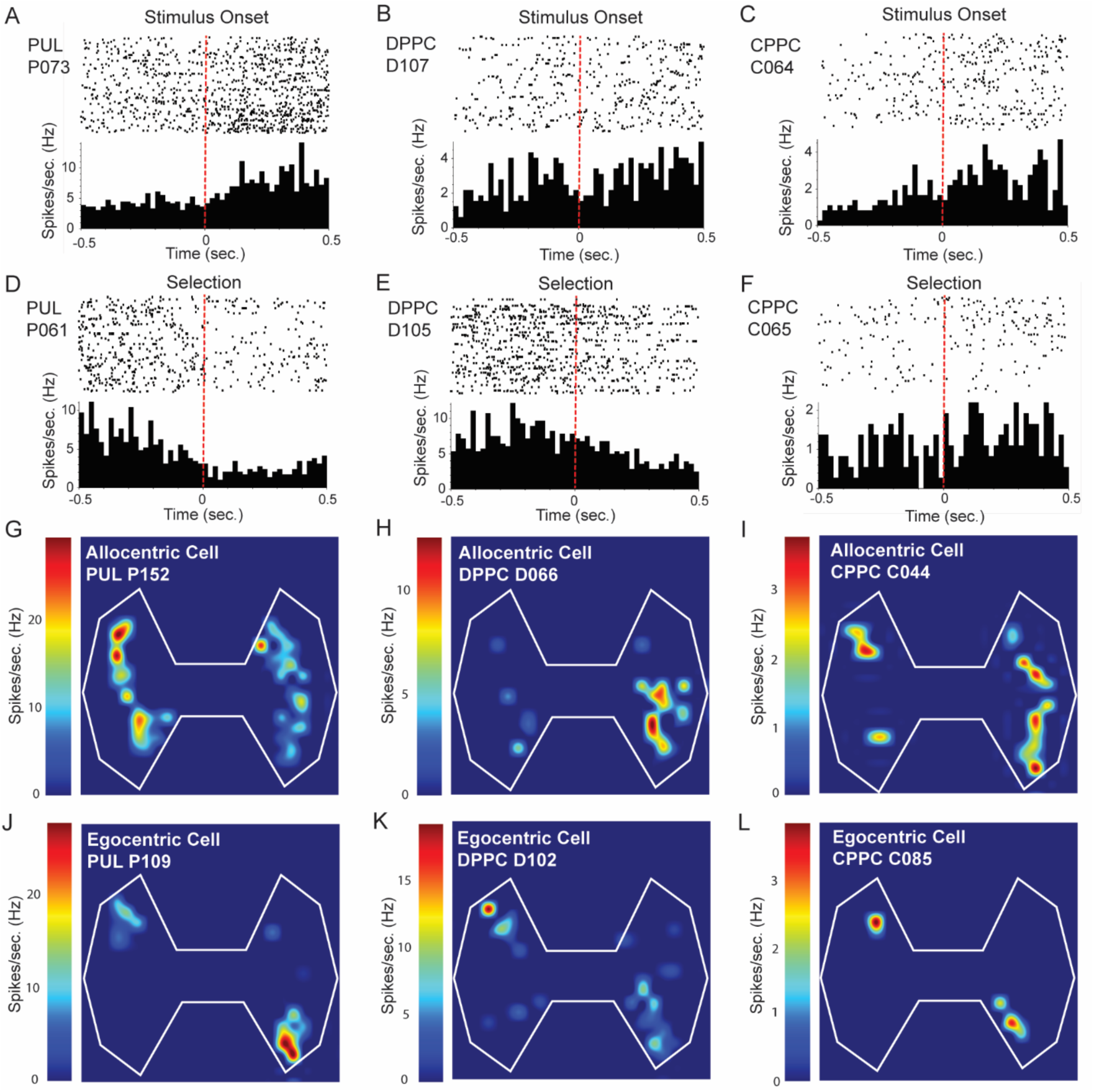
Cells in the pulvinar (PUL), dorsal posterior parietal cortex (DPPC), and caudal posterior parietal cortex (CPPC) signal stimulus onset, selection, and location. (A-C) Perievent raster plots and histograms are shown for example cells from PUL, DPPC, and CPPC that signal stimulus onset (Time 0). (D-F) Perievent raster plots and histograms are shown for example cells from PUL, DPPC, and CPPC that signal selection (Time 0). (G-I) Example cells with allocentric location correlates in the PUL, DPPC, and CPPC. (J-L) Example cells with egocentric location correlates in PUL, DPPC, and CPPC.

For the peri-stimulus and peri-selection epochs, we compared firing rates (FR) of each neuron 500 ms before and after the onset of illumination of the target stimulus (stimulus onset, Figure 1B) and between correct vs incorrect trials (outcome). In each region about one quarter of the cells (∼21% to 29%) exhibited some type of selectivity (Figure 3A, solid bars; Table S1, LED-OFF columns). Overall, the three regions exhibited different patterns of behavioral correlates as represented by the frequencies of cells selective for stimulus, outcome, the conjunction of stimulus and outcome, or that were not selective. For the CPPC the highest proportion of selective cells coded for the conjunction of stimulus onset and outcome followed by stimulus only. The PUL exhibited the opposite pattern. For the DPPC, the highest proportion of selective cells were coded for stimulus only, whereas the proportion coding for the conjunction of stimulus onset and outcome and outcome only were about the same. Chi-square analysis indicated that the patterns of neuronal correlates differed across regions (χ² (6) = 23.863, p = 0.001). Pairwise comparisons indicated that the CPPC was significantly different from the PUL (χ² (3) = 11.063, p = 0.03) and the DPPC (χ² (3) = 14.305, p = 0.009), and the PUL and DPPC were marginally significantly different from each other (χ² (3) = 8.886, p = 0.09).

**Figure 3.**
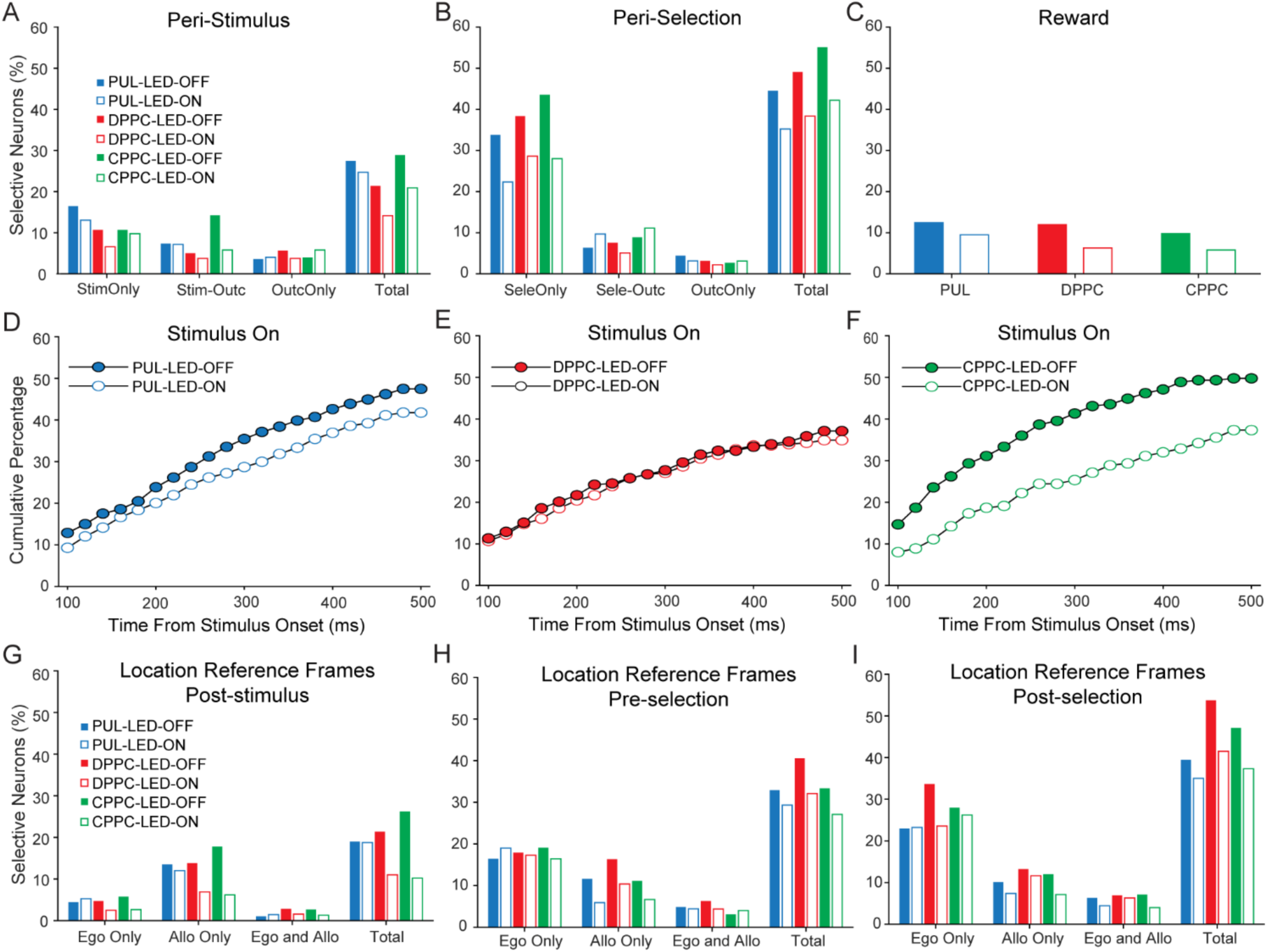
Optical perturbation alters behavioral correlates in the pulvinar (PUL), dorsal posterior parietal cortex (DPPC) and caudal posterior parietal cortex (CPPC). (A-C) Percentage of cells selective for stimulus onset, outcome, or both across the peri-stimulus the peri-selection epochs and for outcome in the reward epoch. (D-F) Time course of emerging selectivity to stimulus offset shown as the cumulative percentage of cells that exhibited a significant difference between pre-stimulus and post-stimulus epochs in 100 ms windows sliding by 20 ms. Lines with solid circles represent LED-OFF trials and empty circles indicate LED-ON trials. (G-I) Percentage of cells selective for egocentric or allocentric locations (east or west) or the conjunction of egocentric and allocentric reference frames during post-stimulus, pre-selection and post-selection epochs.

We were also interested whether there were differences across regions in the latency of neurons to become responsive to stimulus onset. For each neuron, we found the time when its mean FR in the post-stimulus epoch differed from the mean FR during the pre-stimulus epoch. We then constructed a cumulative percentage of cells that were different in two consecutive 100 ms windows sliding by 20 ms (Figure 3D-F, solid markers). Visual inspection of the curves both the PUL and CPPC distributions were higher than the DPPC distribution in control conditions. Moreover, the CPPC distribution appears more curvilinear, such that cells become responsive more rapidly in the CPPC than the DPPC. Using the Kolmogorov-Smirnov nonparametric test for pairwise comparisons, we found that the DPPC was significantly different from the CPPC (Z = 1.897, = 0.003) and marginally significantly different from the PUL (Z = 1.423, p < 0.10). The PUL and CPPC were not different from each other (Z = 0.791, p = 1.00).

To summarize responses to stimulus onset under control conditions, for all three regions the combination of cells signaling stimulus onset and the conjunction of stimulus onset with outcome was greater than cells signaling outcome alone. The CPPC stands out in that the largest proportion of cells signaled the conjunction of stimulus onset with outcome, suggesting that CPPC coding for stimulus onset is predictive of good performance, an interpretation consistent with a role in stimulus driven attention. In terms of the latency to respond to stimulus onset, the CPPC and PUL showed shorter latencies and more responsive cells compared with the DPPC, suggesting the CPPC and PUL respond more rapidly to task relevant stimuli, consistent with a greater role in stimulus-driven attention.

We then analyzed behavioral correlates of target selection. We compared FR 500 ms before the animal’s head entered a target zone (pre-selection) and 500 ms after (post-selection) for correct and incorrect LED-OFF trials (Figure 3B, solid bars; Table S1, LED off columns). The majority of cells in all three regions signaled target selection, and the proportions of cells that signaled target selection, outcome, or both were about the same across regions. Multiway chi-square analysis indicated no differences across the three regions in the frequencies of cells coding for target selection, outcome, the conjunction of section and outcome, or showing no selectivity (χ² (6) = 10.447, p = 0.11). All three regions exhibited a preponderance of coding for target selection along with much smaller proportions of conjunctive selectivity with the fewest cells coding for outcome only. We also analyzed neuronal correlates of trial outcome during the reward-approach epoch, in which the animal had just entered the food area after selecting a target location (Figure 3C, solid bars; Table S1, LED off columns). Across the three regions we observed similar proportions of outcome selective and nonselective cells (χ² (2) = 1.082, p = 0.58) with proportions ranging from ∼10% to ∼12%.

To assess neuronal correlates of egocentric and allocentric location of the target stimulus under control conditions, we pooled trials in which the target stimulus was on the same side of the maze (east vs. west) for allocentric location correlates, and pooled trials in which the target stimulus was at the same egocentric location (left, right, and center) for egocentric location correlates (Figure 3G-I). We analyzed epochs for which spatial location of the target was task-relevant: post-stimulus, pre-selection, and post-selection (Table S2). We were first interested in whether the pattern of correlates (egocentric, allocentric, the conjunction of egocentric and allocentric reference frames, or no selectivity) differed across epochs. We predicted differences because the behavior with respect to location is different across epochs. In the post-stimulus epoch, the animal is in the center of the maze and facing the location where the target appeared, in the pre-selection epoch the animal is approaching the target location, and in the post-selection epoch the animal is in the target location. An examination of the correlates (Figure 3G-I; Table S2) shows that the numbers of selective cells increased with each successive epoch, largely accounted for by cells coding for egocentric locations in pre- and post-selection epochs. Conjunctive coding for egocentric and allocentric locations also increased modestly. This was confirmed by a multiway (correlate by region by epoch) chi-square analysis (χ² (6) = 18.080, p = 0.006). Pairwise comparisons indicated that the post-selection pattern of selectivity differed from both the post-stimulus and pre-selection patterns (χ² (6) = 16.189, p = 0.04 and χ² (6) = 20.396, p = 0.006, respectively), but the post-stimulus and pre-selection patterns were not different from each other (χ² (6) = 7.450, p = 0.72). Unsurprisingly, allocentric-only coding was stable across epochs, possibly because allocentric location is not relevant to making correct choices and is only relevant for initiating the next trial.

Examination of regional differences for each epoch indicated that, in the post-stimulus epoch, the proportions of cells that signaled egocentric, allocentric, the conjunction of egocentric and allocentric reference frames, or were not selective were similar across regions (χ² (6) = 7.450, p = 0.28), nor were there differences in the pre-selection epoch (χ² (6) = 9.377, p = 0.15). However, in the post-selection epoch, when the animal was positioned in a task relevant target location, there were differences across regions (χ² (6) = 17.011, p = 0.009). Specifically, the DPPC exhibited the largest proportion of egocentric correlates followed by the CPPC. This was confirmed by pairwise comparisons indicating a significant difference in reference frame correlates between PUL and DPPC ((χ² (3) = 16.732, p = 0.003) and no significant differences between PUL and CPPC (χ² (3) = 3.735, p = 0.88) or between CPPC and DPPC (χ² (3) = 2.965, p = 1.00).

In summary, our findings during control conditions were consistent for a role in visuospatial attention for all three regions. The CPPC differed from the other regions coding more strongly for the conjunction of stimulus onset possibly suggesting a stronger role in stimulus driven attention. All three regions coded strongly for target selection. All three regions coded for allocentric and egocentric location of the target as well as conjunctions of the two reference frames. Allocentric correlates were stable across epochs which may reflect the lack of task relevance for target selection. Egocentric correlates, however, increased across epochs and were highest in the pre- and post-selection epochs, especially in the DPPC. This may reflect a greater role for the DPPC in action selection.

### Optical suppression of PUL neurons had modest behavioral impact on visuospatial attention

PUL neurons were stimulated optogenetically on half the trials. Interestingly, optical suppression had no impact on behavioral accuracy and resulted in only a modest slowing of latency to make a selection. Accuracy was 65.4 ± 3.6% in LED-OFF trials and 66.1 ± 3.8% in LED-ON trials (F(1) =1.323, p = 0.30). Overall, response latencies were faster on correct trials than incorrect trials and faster on LED-OFF trials than LED-ON trials: LED-OFF, correct, 1.29 ± 0.11 sec, incorrect, 1.73 ± 0.18 sec; LED-ON trials, correct, 1.33 ± 0.14 sec, incorrect, 1.79 ± 0.20 sec. This was confirmed by two-way ANOVA showing a main effect of outcome (F(1) =20.702, p = 0.006) and a marginally significant main effect of LED condition (F(1) =5.711, p = 0.06) with no interaction (F(1) =0.406, p = 0.55). Overall, optical suppression in the PUL did not impair accuracy, but may have slowed latencies to choose. One possibility of this modest impact on behavior may be related to the level of difficulty in this VSA task. Another possibility may be relevant to changes of neuronal coding in these brain regions. To address this, we examined neuronal activity in several aspects in the next series of analyses.

### Neuronal correlates of visuospatial attention reorganized during PUL suppression trials

#### Optical suppression of the PUL decreased overall selectivity

Next, we assessed the effects of perturbing this circuit at the level of PUL with optical suppression. We thus identified neuronal correlates using the same analysis techniques we used during control conditions, using only the trials in which the PUL was optically suppressed. We expected to see a decrease in neuronal correlates of behavior in the PUL and perhaps alterations in the downstream DPPC and CPPC. We found optical suppression decreased selectivity overall in both the peri-stimulus epoch and the peri-selection epoch (χ²(2) = 15.589, p = 0.001 and χ²(2)=8.918, p = 0.01, respectively). The overall decrease in the number of selective cells in LED-ON trials compared with the LED-OFF trials confirmed that optical suppression impacted neuronal correlates of the task, even though performance on the task was largely spared.

#### Optical suppression in the PUL influences responses to stimulus onset in the PUL, DPPC, and CPPC

A primary goal of the study was to understand how neuronal correlates changed with optical suppression of the PUL. Beginning with responses to stimulus onset, we identified cells that were selective only under LED-OFF conditions, were selective only under LED-ON conditions, showed the same correlates under both conditions, changed correlates between conditions, or were nonselective under both conditions. For example, a cell that signaled stimulus onset and outcome in control conditions might change to a stimulus onset only cell under LED-ON conditions. Examination of Figure 4A and Table S3 indicates that, within each region, the largest proportion of cells selective at some point were in the LED-OFF only category, followed LED-ON only category, altered correlates during LED-ON and OFF, and finally the same correlates during both conditions. Compared with the other regions, however, the DPPC exhibited the largest proportion of cells that were selective only under control conditions and the smallest proportion that showed the same correlates under both conditions. A multiway chi-square analysis indicated regional differences in how correlates changed in response to stimulus onset (χ²(8) = 22.137, p = 0.005). Pairwise comparisons indicated that the DPPC was significantly different from the PUL (χ²(4) = 21.131, p = 0.003) and marginally significantly different from the CPPC (χ²(4) = 10.989, p = 0.08). The CPPC and PUL were not different from each other (χ²(4) = 2.713, p = 1.00). Thus, the impact of optical suppression was highly similar across the PUL and CPPC, but different from the DPPC in which a very small number of cells showed the same correlates under both conditions. Importantly, most cells selective in at least one LED condition exhibited altered selectivity, rather than a simple loss of selectivity, providing evidence for functional reorganization in all regions.

**Figure 4.**
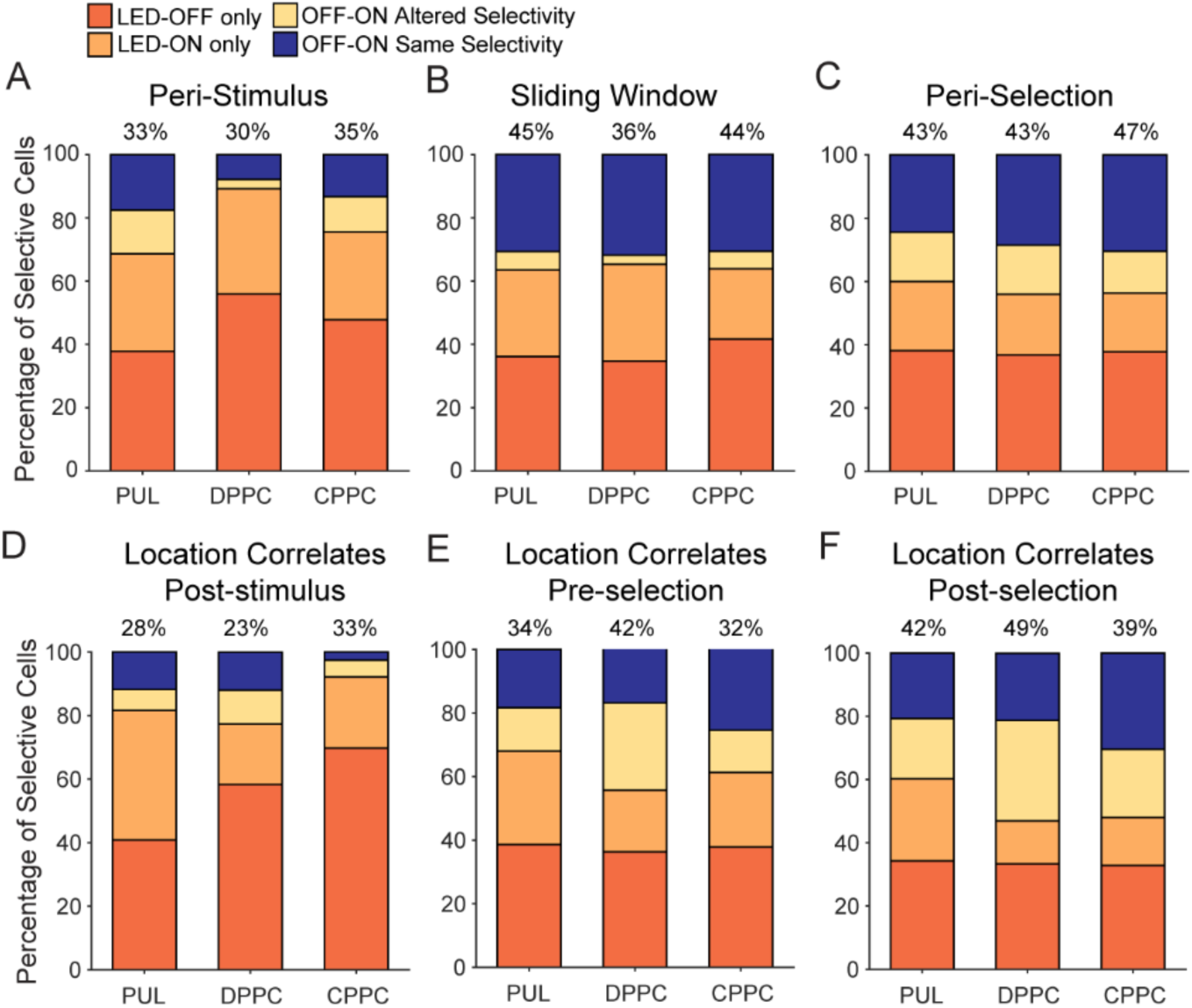
Most selective cells in the pulvinar (PUL), dorsal posterior parietal cortex (DPPC) and caudal posterior parietal cortex (CPPC) show altered behavioral correlates with optical perturbation. (A-B) Impact of optical perturbation on responses to stimulus onset in the Peri-Stimulus and Sliding Window analyses. Above each bar is the percentage of recorded cells (PUL, n = 474; DPPC, n = 318; CPPC, n = 225) showing some type of behavioral correlate during at least one condition. (C) Impact of optical perturbation on correlates of choice behavior in the Peri-Selection epoch. (D-F) Impact of optical perturbation on location correlates during post-stimulus, pre-selection, and post-selection epochs, respectively.

To further understand this reorganization, we used the sliding window analysis to characterize the impact of PUL suppression on the temporal dynamics of neuronal responses to stimulus onset. For each neuron, we found the time in the post-stimulus epoch when its mean FR differed from the mean FR during the pre-stimulus epoch. We then constructed cumulative distributions for each region in the LED-OFF and LED-ON conditions resulting in 6 distributions (Figure 3D-F; Figure S1). Visual inspection of the curves suggests that the CPPC was most impacted by optical suppression of the PUL and the DPPC was least impacted. Using the nonparametric Friedman test for repeated measures, we found overall regional differences across the LED-OFF vs LED-ON distributions (χ²(5) = 88.625, p = 0.001). A Kruskal-Wallis test indicated that central tendencies differed across the three regions in the LED-OFF condition (H(2) = 10.930, p = 0.008), but not the LED-ON condition (H(2) = 1.173, p = 1.00). Using the Kolmogorov-Smirnov nonparametric test for pairwise comparisons we tested whether each region was different across the LED-OFF and LED-ON conditions. We found no difference across conditions for the PUL (Z = 0.791, p = 1.00) or the DPPC (Z = 0.316, p < 1.00), but optical suppression significantly reduced the number of responsive cells and retarded the time from stimulus onset to responsivity for cells in the CPPC (Z = - 3.585, p < 0.003). In summary, optical suppression appeared to reduce the diversity of temporal responses across regions, and temporal dynamics in the CPPC were most affected by optical suppression. This suggests the CPPC relies more heavily on PUL than the DPPC in the signaling of task-relevant stimuli.

#### Optical suppression in the PUL influences PUL, DPPC, and CPPC cells during choice behavior

We were also interested in whether correlates changed with optical suppression during decision making or choice behavior. Accordingly, for the peri-selection epoch we again identified and categorized cells as selective only under LED-OFF conditions, selective only under LED-ON conditions, showing the same correlates under both conditions, showing different correlates between conditions, or nonselective under both conditions (Figure 4C; Table S3). Proportions of cells in each category were remarkably similar across the three regions. In all three regions, the largest category were cells selective only during LED-OFF trials (∼37-38%) followed by cells that showed the same correlates in both conditions (∼24-30%). Next were cells that were selective only during LED-ON trials (∼19-22%) followed by cells that showed altered correlates across LED-ON and LED-OFF conditions (∼13-16%). This was confirmed by multiway Chi-square analysis indicating a lack of regional differences in the patterns of changes in selectivity (χ²(8) = 9.241, p = 0.32).

Neuronal correlates of trial outcome during the reward epoch showed little impact of optical suppression (Figure 3C; Table S1). Across the three numbers of cells that were outcome selective or not selective under control and LED-On conditions were not different as confirmed by multiway chi-square analysis (χ² (2) = 3.892, p = 0.14).

In summary, optical suppression impacted all three regions in roughly the same way. Notably, during the peri-selection epoch in all three regions most selective cells showed altered behavioral correlates with optical suppression of the PUL, ranging from ∼70% to ∼76% (Figure 4C).

#### Optical suppression in the PUL influences neuronal coding of location reference frames in the PUL, DPPC, and CPPC

We next examined how correlates of allocentric and egocentric reference frames changed with optical suppression. For both LED-OFF conditions and LED-ON conditions, we identified cells that were selective for egocentric, allocentric, or the conjunction of egocentric and allocentric reference frames (Figure 3G-I; Table S2) and cells that were never selective. We then binned cells into five categories: selective only under LED-OFF conditions, selective only under LED-ON conditions, same correlates under both conditions, changed correlates between conditions, or no correlates under either condition (Figure 4D-F; Table S3). We categorized cells separately for the post-stimulus, pre-selection, and post-selection epochs.

We first examined whether the impact of optical suppression on location correlates differed across epochs. We predicted differences because the behavior with respect to location is different across epochs. An examination of Table S2 shows that the proportions of selective cells increased with each successive epoch for all three regions and in both LED-OFF and LED-ON conditions. This was confirmed by a multiway (led impact by region by epoch) chi-square analysis (χ² (8) = 55.608, p = 0.001). Pairwise comparisons indicated the impact of optical suppression differed across each pair of epochs: post-stimulus vs pre-selection, χ² (8) = 33.781, p = 0.003; post-stimulus vs post-selection, χ² (8) = 39.177, p = 0.003; and pre-selection vs post-selection, χ² (8) = 50.495, p = 0.003.

Further examination of Figure 4D-F and Table S3 suggests regional differences within each epoch. In the post-stimulus epoch, the PUL showed roughly equal proportions of LED-On only and LED-Off only cells, whereas both the CPPD and DPPC had the highest proportion of cells in the LED-OFF only category. Paired comparisons confirmed that the PUL differed from the DPPC (χ² (4) = 14.763, p = 0.02) and the CPPC (χ² (4) = 18.958, p = 0.003), and the DPPC and CPPC were marginally different from each other (χ² (4) = 10.693, p = 0.09). For the pre-selection epoch, the PUL and CPPC have similar proportions across categories, but the DPPC showed a greater proportion of cells that did not change with suppression. The PUL differed from the DPPC (χ² (4) = 18.501, p = 0.003) but not the CPPC (χ² (4) = 2.818, p = 1.00), and the DPPC and CPPC were marginally different from each other (χ² (4) = 10.513, p = 0.10). For the post-selection epoch, again, the PUL was similar to the CPPC, but not the DPPC. The PUL differed from the DPPC (χ² (4) = 22.494, p = 0.001) but not from the CPPC (χ² (4) = 8.398, p = 0.23), and the DPPC and CPPC were marginally different from each other (χ² (4) = 7.956, p = 0.28).

In summary, the way in which optical perturbation impacted regional selectivity differed across behavioral epochs and across regions. The proportion of cells that were selective in some category increased across epochs (Table S3). However, the proportion of selective cells that reorganized decreased across epochs (Table S3, right column), such that reorganization was strongest in the post-stimulus epoch in which subjects are in the center of the maze at the appearance of the target stimulus. Interestingly, regional differences in the patterns of LED impact were also strongest in the post-stimulus epoch. Overall, these results are consistent with the notion that optical perturbation of the PUL has more impact over stimulus-driven attention that controlled attention or action selection.

#### Spatiotemporal redistribution of location information after optical perturbation

In the sections above we used spike rate analysis to categorize isolated single units or neurons as responsive or not to a particular task relevant event such as onset of a stimulus in a target location or the selection of a target location. Using that method, we have shown that individual neurons in the PUL, DPPC, and CPPC robustly encode behaviorally relevant information throughout the VSA task and that the proportion of neurons coding for task relevant information, as well as any given neuron’s particular behavioral correlates, were influenced by optical suppression of PUL (Figure 3), even though there was no behavioral impairment in the task.

These results raised two additional questions about the effects of optical suppression that could be addressed using encoding and decoding analyses. First, how does optical suppression in the PUL change encoding of multiple VSA task variables in local (PUL) and distant (DPPC and CPPC) areas? Second, given that there was no impact of suppression on accuracy in the task, how does location information represented by neural ensembles in these regions support visuospatial attention during optical suppression?

To characterize how task information was encoded in population level neural activity, we used multiple linear regression models to determine which variables of the task were encoded by each neuron during five different 500-ms epochs: pre-stimulus, post-stimulus, pre-selection, post-selection, and reward (see Methods). Trial-wise FR in each epoch were predicted by the following: allocentric information, egocentric information, outcome, and LED ON/OFF (Equation 1). We found that the proportion of neurons encoding allocentric, egocentric, and outcome information gradually increased across epochs (Figure 5A, 5B, and 5C, left panels). However, less than 5% of neurons from all three areas encoded LED-OFF versus LED-ON, suggesting little effect of optical suppression on neural encoding (Figure 5A, 5B, and 5C, purple bars in left panels). This is inconsistent, however, with our earlier result that found differences in selective cells on LED-ON trials during the peri-stimulus and peri-selection epochs. To understand the subtle effects of optical suppression, we reran the regression analysis separately in two groups of trials, LED-OFF and LED-ON trials (Equation 2). Comparing results between two groups revealed that almost all selectivity for all three predictors was greater in LED-OFF trials compared to LED-ON trials (Figure 5A, 5B, and 5C, middle and right panels). Therefore, optical suppression decreased encoding of location and outcome information in three brain areas across epochs, although some encoding of all three predictors remained during LED-ON.

**Figure 5.**
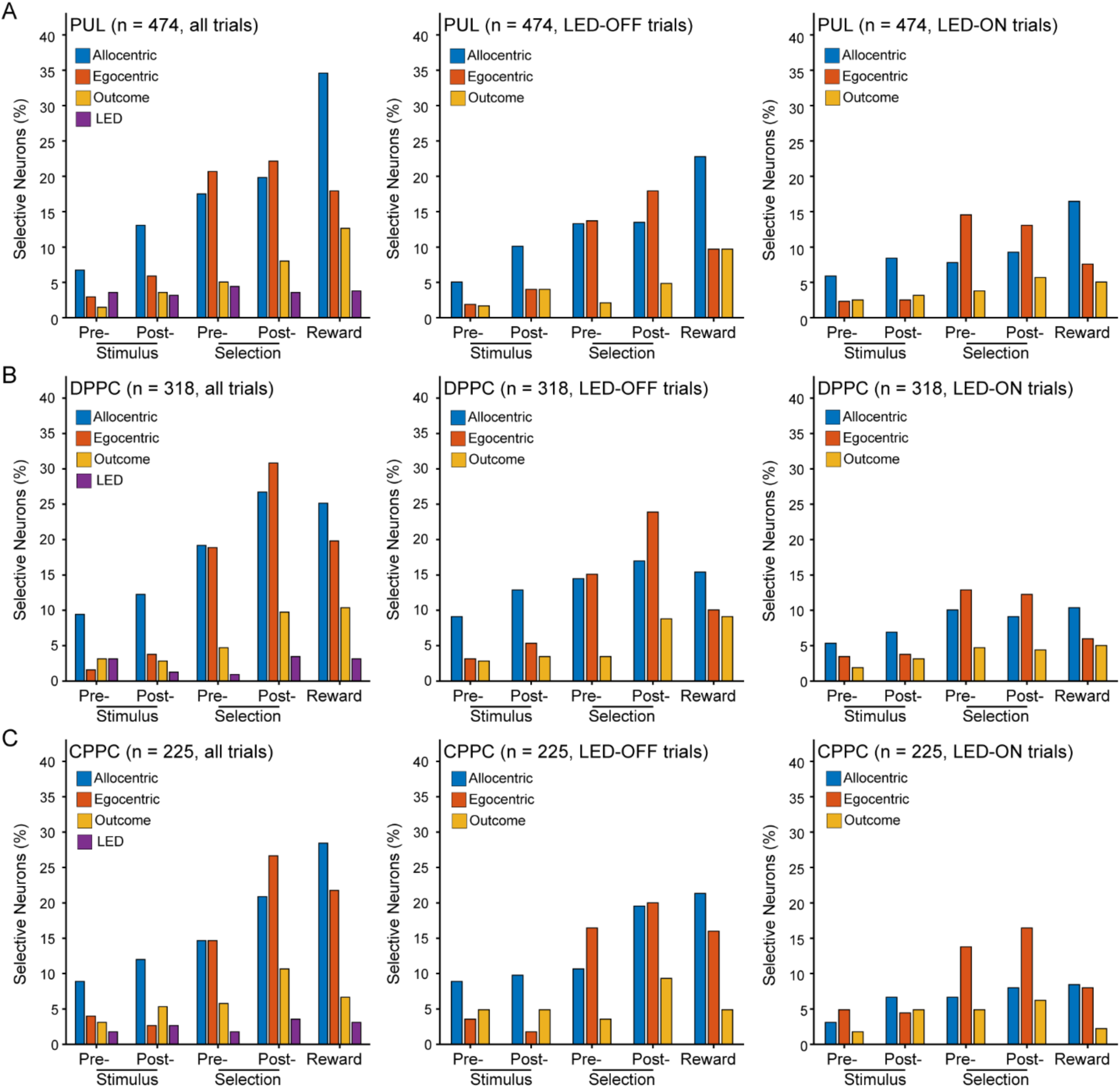
Regression analysis shows that cells in the pulvinar (PUL), dorsal posterior parietal cortex (DPPC) and caudal posterior parietal cortex (CPPC) are selective for stimulus, selection, reward and location. Percentages of selective neurons in the PUL (A), DPPC (B) and CPPC (C) are determined by multiple linear regressions. Left panels show results from all trials (using both LED ON and OFF) including information of allocentric location, egocentric location, outcome and LED. The middle and right panels are the reduced model regressions which included information of allocentric location, egocentric location, and outcome, run separately on LED-OFF (middle) and LED-ON (right) trials. Each panel shows five different 500-ms epochs are on the X-axis.

To examine whether the time courses of neural encoding changed with optical suppression, we calculated the coefficient of partial determination (CPD) in sliding windows before and after stimulus onset or selection and sorted neurons according to encoding latency (Figure 6 and Figure S2). We did this separately for three predictors (allocentric location, egocentric location, and outcome) in each brain area, on LED-OFF and LED-ON trials. In Figure 6A, the left panel shows the time course of encoding allocentric information in PUL before and after selection during LED-OFF trials. The superimposed blue curve shows the encoding latencies for each neuron. The middle panel shows encoding among the same group of PUL cells except on LED-ON trials, where the neurons were sorted in the order determined by the LED-OFF trials in the left panel. In this case, the latency curve was no longer clearly visible, suggesting that the temporal order of neuron encoding was not preserved. However, the allocentric information was still encoded during LED-ON trials, as the response latency curve was clearly visible if the same LED-ON data were re-sorted according to the latencies observed on the LED-ON trials (Figure 6A, right panel). For visualization, a new latency curve for these trials was superimposed in red.

**Figure 6.**
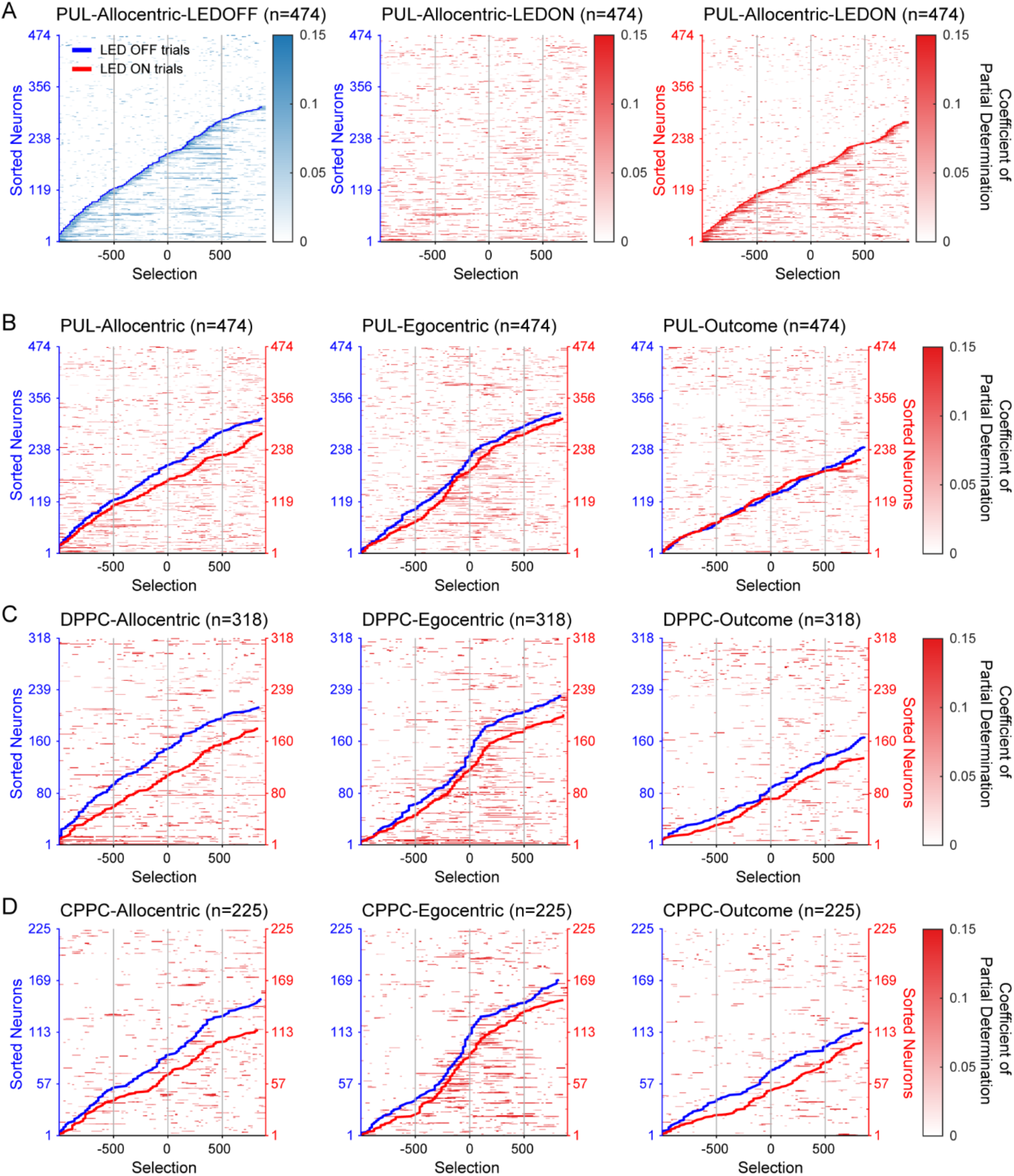
Regression analysis shows that information encoding in populations is redistributed with optical perturbation. (A) Plots of allocentric information encoding in the pulvinar (PUL) aligned to the time of selection, illustrating how latency curves computed from CPDs were compared. Encoding latency was defined as when the first of three consecutive sliding windows exceeded the criterion of >= 95th percentile of 1000 permutation tests on each time window. The left panel shows neurons sorted according to encoding latency on LED-OFF trials, the right panel shows the same neurons sorted on LED-ON trials. The middle panel shows LED-ON trials sorted according to the order established in LED-OFF trials. Blue and red lines were superimposed on the heatplots for visualization. (B-D) Blue and red lines show latency curves from sorted LED-OFF and ON trials, superimposed on the heatmap pattern of LED-ON trials sorted according to the LED-OFF order. These integrated plots show encoding of three predictors, allocentric information (left), egocentric information (middle), and outcome (right) in the (B) PUL, (C) dorsal posterior parietal cortex (DPPC), and (D) caudal posterior parietal cortex (CPPC).

Together, these data suggest that optical suppression of the PUL reorganized the temporal order of neuron encoding in PUL but did not cause the full loss of allocentric, egocentric, or outcome information. Importantly, the subgroup of neurons that formed the blue latency curve in the left panel is not necessarily the same subgroup of neurons forming the red latency curve, as shown in Figure 6, as only neurons whose encoding reached significance were included in each condition. We applied the same methods to all three task variables and three areas (Figure 6B-D and Figure S2). The remaining panels of Figure 6 show a condensed representation of the analysis, in which we extracted the latency curves from LED-OFF (blue) and LED-ON (red) trials (as in left and right panels of Figure 6A), and superimposed them on the CPD heatmap patterns when LED-ON trials were sorted according to the order determined by LED-OFF trials (as in the middle panel of Figure 6A; color bar in red). In general, we found similar effects across brain regions, where latency curves were comparable in LED-OFF and LED-ON conditions, with slightly higher encoding during LED-OFF, but optical suppression disorganized time courses, potentially by another subgroup of neurons, in LED-ON trials, as the background heatmaps don’t clearly follow the red or blue curves.

Together, these results show that optical suppression changed the neural encodings of location information in PUL, DPPC, and CPPC. However, instead of general suppression of encoding, there was a re-organization of task information within the population, resulting in information encoded with different time courses and in different subgroups of neurons. Given this, we next explored how location information was represented by neural ensembles with or without optical perturbation.

#### Functional reorganization of location information with optical perturbation

To explore how neural ensembles represented location information, we created pseudo-populations of recorded units from each of the three areas and separated them into LED-OFF and LED-ON trials. We then used these populations to decode each of six different locations in which the target could appear (see Methods). To assess the similarity of the representation across LED-OFF and LED-ON trials, we assessed the following TRAIN and TEST conditions: 1) train on LED-OFF trials and test on held-out LED-OFF trials; 2) train on LED-ON trials and test on held-out LED-ON trials; and 3) train on LED-OFF trials and test on LED-ON trials. Based on the sorted latency curves in the previous analysis (Figure 6), we expected the six locations to be represented by neural ensembles both with and without optical perturbation, but that optical perturbation may cause the information to be redistributed within the ensemble, so that cross-condition decoding performance would be low.

During LED-OFF trials, we found that location information could be decoded from ensembles of PUL, DPPC, or CPPC neurons (Figure 7 and Figure S3). Figure 7A-C, first row, are shown by posterior probabilities, which were highest for each of the six correct locations, depicted on the diagonal of each plot (chance decoding = 16.7%). Therefore, neural ensembles in PUL, DPPC, and CPPC have unique firing patterns to represent each location in the VSA task. During LED-ON trials, location information was clearly represented by PUL, but overall decoding was slightly lower in DPPC and CPPC compared to LED-OFF trials (Figure 7A-C, second row). In particular, posterior probabilities for the middle location on west trials were lower than chance. Therefore, although allocentric and egocentric information was represented by single neurons in DPPC and CPPC in the previous analysis (Figure 6), individual locations were not as distinguishable based on population firing patterns. Nonetheless, 5 of 6 locations were decodable above chance during LED-ON trials, demonstrating that considerable target information is preserved with optical suppression of PUL.

**Figure 7.**
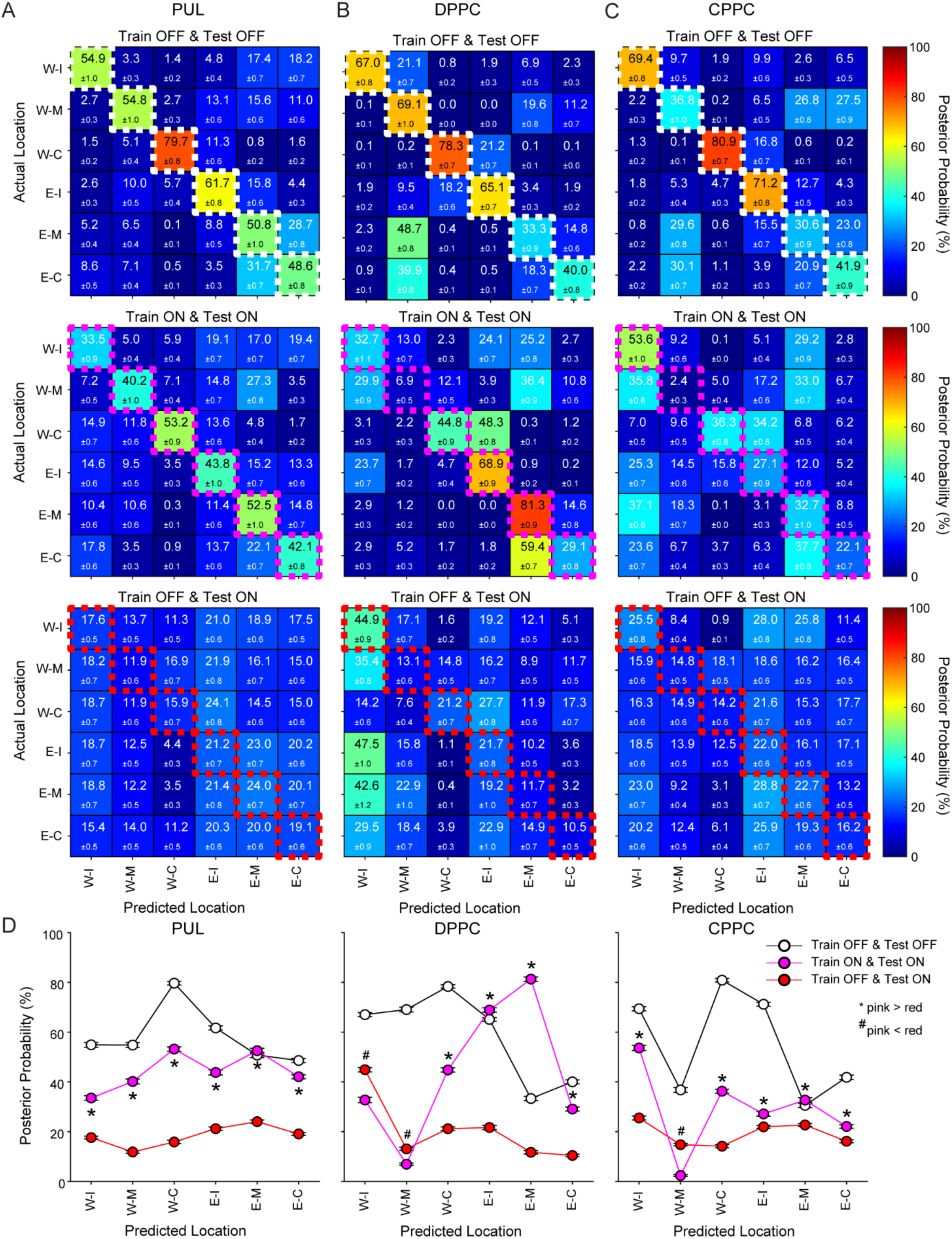
Cross-condition decoding analysis also shows that information in ensemble is redistributed with optical perturbation. (A-D) Decoded location information for the post-selection epoch is shown for the pulvinar (PUL), dorsal posterior parietal cortex (DPPC) and caudal posterior parietal cortex (CPPC). Decoded location information for the other epochs is shown in Figure 8 and Figure S2. (A-C) Columns show decoding results from different brain areas, and the first three rows show different Train & Test conditions. The first row shows Train OFF & Test OFF, in which decoders were trained on LED-OFF trials and tested on held-out LED-OFF trials. The second row shows Train ON & Test ON, where decoders were trained on LED-ON trials and tested on held-out LED-ON trials. The third row shows Train OFF & Test ON, where the decoder was trained on LED-OFF trials and tested on LED-ON trials. Each 6-by-6 heatmap shows the actual location on the y-axis, with the posterior probabilities of all 6 possible locations, so that correct locations are on the main diagonal (outlined). Each element in heatmaps is the mean posterior probability for each location ± sem. ‘W’ = stimulus turns on in west trials, ‘E’ = stimulus turns on in east trials. ‘I’ = ipsilateral, ‘M’ = middle, and ‘C’ = contralateral location relative to the recorded hemisphere. (D) Posterior probabilities of correct predictions (diagonal) from the three conditions above are reproduced. * = Train ON & Test ON significantly higher than Train OFF & Test ON on post-hoc comparisons (p < 0.001). ^#^ = Train OFF & Test ON significantly higher than Train ON & Test ON on post-hoc comparisons (p < 0.001). Bonferroni corrected for multiple comparisons.

Finally, we trained decoders with the neural ensembles in the LED-OFF trials and tested them with the same ensembles in the LED-ON trials (Figure 7A-C, third row). Here, decoding was at or below chance levels in all locations and brain areas, except for the ipsilateral location in the west trials in DPPC. Statistical tests revealed that nearly all comparisons of LED-OFF, LED-ON, and cross-condition decoding were significant, with lowest performance on the train OFF & test ON condition (Figure 7D) (Two-way ANOVA of locations x decoding conditions on posterior probabilities: PUL, locations F_5,17982_ = 132.75, p < 0.001, conditions F_2,17982_ = 3554.29, p < 0.001, interaction F_10, 17982_ = 79.23, p < 0.001; DPPC, locations F_5,17982_ = 526.43, p < 0.001, conditions F_2,17982_ = 3510.38, p < 0.001, interaction F_10, 17982_ = 702.77, p < 0.001; CPPC, locations F_5,17982_ = 712.89, p < 0.001, conditions F_2,17982_ = 3451.48, p < 0.001, interaction: F_10, 17982_ = 276.56, p < 0.001). This shows that the specific firing patterns that represent locations during LED-OFF trials are distinct from those in LED-ON trials, and the information that was preserved in LED-ON trials was not simply a subset of the information encoded during LED-OFF trials.

Post-hoc tests compared decoding performance, quantified by the mean posterior probabilities of all correct locations, across the three training and testing conditions (Figure 8). During the pre-stimulus epoch, decoding in all conditions was close to the chance levels (16.6%) (Figure 8A). After stimulus onset, decoding in the train OFF & test OFF condition increased (Figure 8B-D), while decoding performance in the train ON & test ON condition only increased during the post-selection epoch, when the rat had physically moved to the decoded location (Figure 8D). Decoding performance in the train OFF & test ON condition was consistently low across all epochs (Figure 8, red bars in all panels). The poor cross-condition decoding is further evidence that represented location information in the train OFF & test OFF condition was reorganized after optical perturbation. This was true even in downstream areas such as DPPC and CPPC that did not receive optical perturbation directly. The fact that location information was only decoded in the train ON & test ON condition after the rat had locomoted to the selected area suggests that physical location, rather than stimulus information, might be forming the location representations in the presence of optical perturbation.

**Figure 8.**
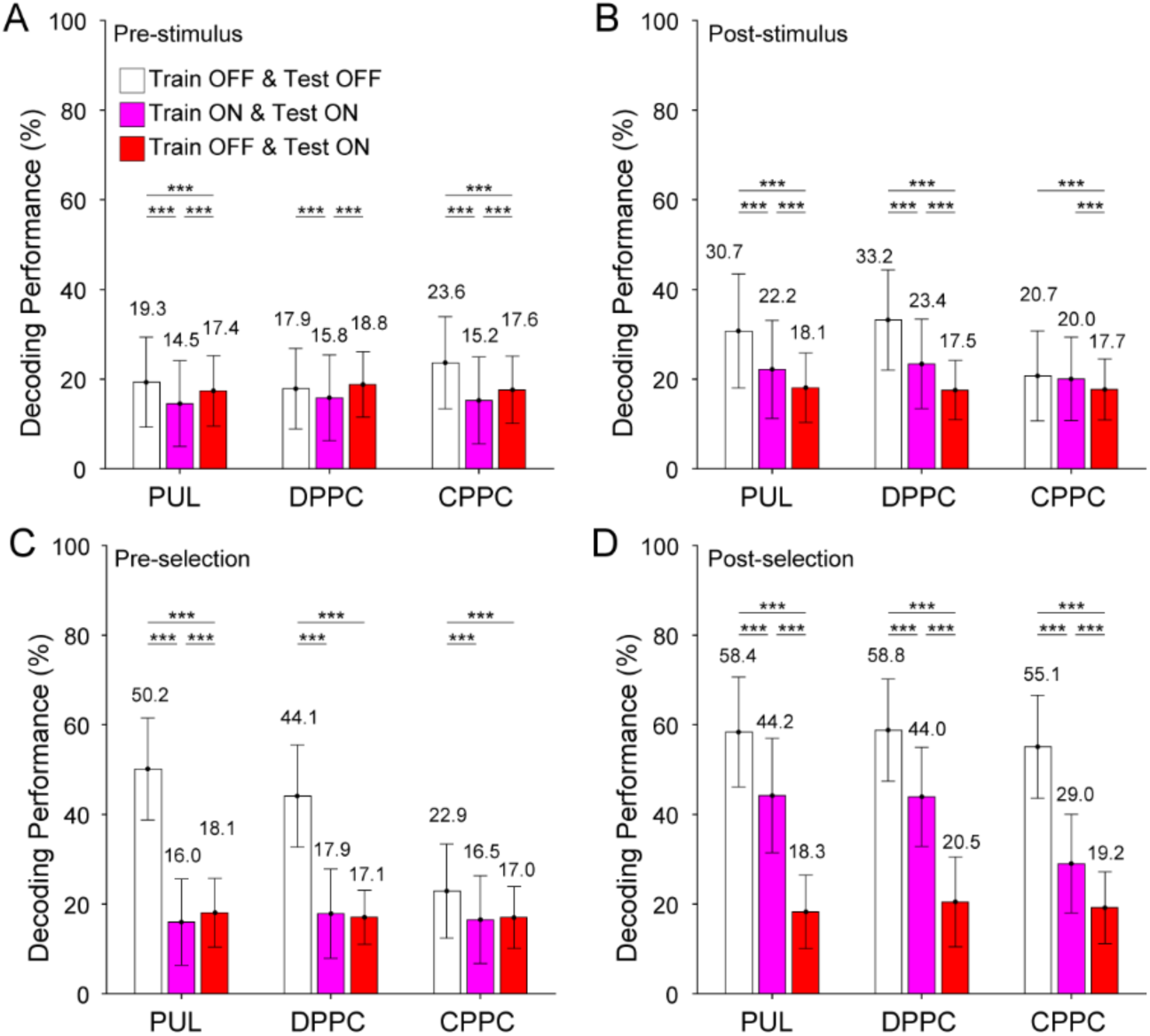
For the same LED conditions (OFF-OFF and ON-ON), decoding performance improves across trial epochs. (A-D) Decoding performance, defined as mean posterior probabilities from six cue locations, is shown separated by brain area, condition, and epoch. (A) Decoders in three Train-Test conditions from the pulvinar (PUL), dorsal posterior parietal cortex (DPPC) and caudal posterior parietal cortex (CPPC) showed similarly low accuracy during the pre-stimulus epoch (chance = 16.7%). (B-D) After stimulus onset, decoding in the train OFF & test OFF condition began to increase, while decoding in the train ON & test ON condition only increased during the post-selection epoch. Decoding accuracy in the train OFF & test ON condition was consistently low across epochs. Error bars indicated standard deviation. *** p < 0.001.

Finally, to take a closer look at firing patterns in the LED-OFF and LED-ON trials, we projected firing patterns into a subspace composed of the first 5 principal components, and found that, across trials, activity in PUL, DPPC and CPPC occupied partially orthogonal subspaces, consistent with the notion that there were distinct firing patterns with and without optical perturbation (Figure S4). In practice, this temporally stable subspace for representing target locations during the optical perturbation likely allowed the animals to perform the task with no major behavioral changes, since it allowed for the readout of location information encoded during the optical perturbation, or after distractor interference ^29^.

## Discussion

In this study, we examined the role of the PUL projections to the DPPC and VPPC in visuospatial attention. We hypothesized that PUL input to the DPPC and CPPC enhances visuospatial information processing and improves visuospatial attention. We predicted that PUL suppression would result in degraded representations of task relevant stimuli in the DPPC and CPPC and that the CPPC might be more impacted given its more prominent role in stimulus-driven attention. We found that PUL suppression did result in lower proportions of selective cells in all three regions. However, many task-related responses were still recovered from all three regions. Analysis of single units and ensembles in all three regions showed that neural activity was functionally reorganized in the presence of optical suppression to support accurate performance in our visuospatial attention task. Additionally, in many analyses, the CPPC was more impacted than the DPPC as we had predicted or the impact on the PUL, itself, and the CPPC were similar, supporting the view that the PUL input to the CPPC supports stimulus-driven attention.

As in our prior studies, we used a bowtie shaped maze allowing us to analyze correlates of top-down and bottom-up attention as well as correlates of allocentric and egocentric frames of reference ^4,12^. In the present study, we recorded neuronal activity in the PUL, DPPC and CPPC simultaneously when animals were under attentional demands as they performed the VSA task. During recording sessions, we used optical perturbation to suppress PUL activity in some trials. Control trials without optical suppression permitted examination of the impact of PUL perturbation on neuronal activity and behavior. Analysis of trials without optical perturbation showed that the PUL and each subregion of the PPC exhibited some differences and some similarities in response to different behavioral phases of the VSA task. First, all three brain regions signaled stimulus onset. In addition, compared to the DPPC, the CPPC had a relatively larger proportion of cells showing stimulus relevant selectivity. Second, cells in the PUL, DPPC and CPPC responded during selection behavior and in the reward epoch. No significant differences among the three brain areas were observed in selection correlates. Third, cells displaying differential firing patterns to task-relevant locations were found in all three brain areas. Fourth, all three regions represented egocentric and allocentric locations, but the proportions of egocentric cells increased as the trial progressed, especially in the DPPC. Finally, we found that over time after stimulus onset, the proportion of selective cells in the PUL and CPPC compared with those in the DPPC increased more rapidly and were significantly higher after stimulus onset. These results show that all three regions represent task-relevant information during the VSA task suggesting that all are involved in stimulus-driven and controlled attention. However, there were also indications that the PUL and CPPC have a greater role in stimulus-driven attention compared with the DPPC.

Results from analyzing LED-OFF trials successfully replicated and extended our previous studies, confirming the roles of the PUL, DPPC and CPPC in a circuit supporting visuospatial attention^4,12,20^. We found that the CPPC was more responsive to stimulus onset than the DPPC, providing evidence that CPPC is more involved in bottom-up attention than DPPC. This is consistent with evidence in humans that ventral PPC is involved in bottom-up attention, reviewed in ^25,30^. Similar to the primate brain, in the rodent brain the CPPC is positioned more ventrally and caudally than the DPPC. The analyses of allocentric location and egocentric location correlates across the three brain areas were consistent with previous findings showing cells in PUL and DPPC respond to task-relevant spatial frames ^4,12,19,20^. However, in the present study, we additionally demonstrated task-relevant location correlates in the CPPC. These results support the view that the PUL, DPPC and CPPC engage heavily in the visuospatial attention network and further provide a strong neuronal basis for the following examination of thalamocortical interaction by suppressing PUL.

By comparing neuronal activity of PUL, DPPC, and CPPC neurons during trials with and without optical perturbation of the PUL, we examined the role of the PUL in an attentional circuit including the DPPC and CPPC. Interestingly, although behavioral changes were not found during the PUL suppression, neuronal activity was altered at both the single-unit and ensemble levels. Overall, at the single-unit level cells either maintained selectivity, changed selectivity, lost selectivity, or became selective, suggesting a functional reorganization of visual information, which may explain the intact visuospatial attention behavior during optical perturbation trials ^31^. First, for the stimulus onset, proportions of selective neurons in DPPC and CPPC significantly decreased during PUL suppression. Over 30% of cells in all three brain regions showed functional changes in behavioral selectivity. We also observed that proportions of selective neurons that responded to animals’ selection behavior in all three areas significantly decreased during PUL suppression. Further, during the selection epoch, over 40% of cells in these brain regions had functional changes in behavioral selectivity. Some cells that responded to allocentric and egocentric locations also changed their location correlates. Second, from the sliding window analysis, PUL inactivation impacted the time course of selectivity after stimulus onset in all three brain areas. Thus, cells in these areas not only changed their functional selectivity, the latency to selectivity to the stimulus onset changed as well. Third, the results from regression models showed that optical perturbation influenced the neural encoding of location information in PUL, DPPC, and CPPC. Instead of general suppression of encoding, follow-up CPD analyses illustrated that re-distribution of task information was observed within the population, resulting in information encoded with different time courses and in different subgroups of neurons. Finally, the results from decoding approaches confirmed that location representations could be decoded by distinct firing patterns with or without optical perturbation. The poor cross-condition, train OFF & test ON, decoding, however, further demonstrated reorganization of task information among neurons in each region. This same pattern was observed in DPPC and CPPC, which did not receive optical perturbation directly but are downstream of PUL. This suggests that the reorganized PUL inputs drove new response patterns in these regions, either directly as different neurons received task-relevant information that differentially drove their responses, or indirectly as these regions reorganized on their own in order to accommodate the new condition. In either case, the reorganized location representations occupied different subspaces, which would allow downstream areas to read out location information in a simple linear fashion ^32^.

The proportions of cells in the PUL and parietal areas that changed their functional correlates during PUL perturbation supports our interpretation that PUL suppression caused a reorganization of information in all three regions. Some of these cells lost their original behavioral correlates while some other cells gained functional selectivity due to the PUL suppression. Surprisingly, despite these changes in neural coding, optical suppression did not impact behavioral accuracy and only modestly impacted latencies to make a choice. These results are similar to the findings in human patients with pulvinar damage. Although the studies are rare, a severe impact on attention is not consistently observed in these patients ^15,33^. The remaining puzzle is how the neuronal changes in the PUL influence the parietal areas and further impact operation of the visuospatial attention network to preserve relevant function. Our results from both encoding and decoding approaches provide an integrated explanation to better understand how task-relevant information is dynamically represented in the brain ^32,34^. At the single-unit level, selectivity was found to shift to different neurons, at different times after stimulus presentation. In ensembles, location information was represented by distinct firing patterns with or without PUL suppression. This reorganization at the ensemble level somewhat preserved decodability of location information in the PUL and PPC areas when the PUL was suppressed, and this may have allowed the attentional circuit to continue to function without noticeable behavioral effects. These results suggest that the task-relevant information can be represented in different ways in the PUL-parietal pathway with minimal or no impact on overt behavior ^29^.

Our results indicate that this functional reorganization of visuospatial information during PUL suppression, locally and its connected PPC subareas, occurs rapidly, raising the question of how this process is orchestrated. One possible mechanism is through interactions in the thalamocortical connections. During the LED-OFF condition, the PPC areas receive direct sensory inputs from the PUL and visual areas ^6,7,13,17^, and bilateral PPC areas exhibit robust homotypic connections ^7^. The reciprocal connections between PUL and PPC areas might provide a route for the PPC to send top-down information back to the PUL for integration and further processing ^4^. When one side of the PUL was suppressed, although the ipsilateral PPC areas lost direct thalamic input due to the PUL inactivation, these PPC areas still received direct visual inputs from other visual areas, such as V1, secondary visual cortex, and the postrhinal cortex ^35,36^. Since the contralateral PPC still received intact thalamic input and could send the information to the ipsilateral PPC, this might allow functional reorganization within the PPC areas. Once the neuronal ensembles of the ipsilateral PPC areas partially reorganize, they could send feedback input to the PUL to rescue partial function. This thalamocortical loop could support the observed the reorganization of neuronal ensembles across three areas during the PUL LED-ON trials and also spare behavioral performance. The robustness of the thalamocortical loop may also explain how patients with pulvinar damage can have spared attentional function.

A small number of non-human primate studies have suggested a role for the pulvinar in coordinating neuronal activity across cortical areas ^37–39^. Prior to the present study, however, evidence for these thalamocortical influences of the attentional network in rodents had been lacking. Here, we provide the first evidence to support the role of the rodent PUL in the attentional modulation of cortical areas in the attention network, specifically the PPC. Further studies are needed to address more detailed neural mechanisms and functional dynamic changes of thalamocortical interactions during visuospatial attention.

## Methods

### Subjects

Subjects were six male Long-Evan rats (Charles Rivers Laboratories, Wilmington, MA). They were individually housed in a temperature-regulated colony maintained on a 12:12 h light:dark cycle. Experiments were carried out in the light phase. All procedures followed the NIH guidelines and were approved by the Brown University animal care and use committee.

### Apparatus

Our task uses the Floor Projection Maze, an apparatus that exploits the natural tendency of rats to attend to items located on or close to the ground and permits automated control over visual stimuli ^27,28^. Essentially, the Floor Projection Maze is a horizontal rear projection screen positioned to allow back-projection of visual stimuli and to serve as a floor to any shaped arena (Figure 1A). The apparatus has a clear Plexiglas subfloor (147.3 cm x 111.8 cm) with a (1.25 cm thick) covered by Dual Vision Fabric (Da-Lite Screen Company, Warsaw, IN), a unity-gain flexible fabric designed for rear screen projection. A thin sheet (Plexiglas 0.6 cm) covers the fabric for protection. Visual stimuli were projected onto the unity gain fabric from below the subfloor using an LCD projector (WT610 projector, NEC Corporation). In this experiment, the enclosure was a bowtie shaped region for presentation of stimuli. Food reward (milk with various flavors) was delivered by two automated pumps (Med Associates, Inc, St. Albans, VT) to stainless steel food ports located at middle region of the maze. Auditory stimuli were controlled by an automated auditory stimulus generator (ANL926, Med Associates, Inc.) and delivered through a speaker located above the maze.

The Floor Projection Maze was interfaced with three Windows PC systems, for location tracking, behavioral control, and neuronal data acquisition. A CinePlex Behavioral Research System (CinePlex Studio and CinePlex Editor) v3.4.1 with a single camera and two additional CinePlex modules, CinePlex Tracking and CinePlex Basic Behavior was used for tracking animal’s location during behavioral performance. The position of the rat was recorded by tracking the centroid of the contour of the rat using the CinePlex Tracking module. Position data are analyzed online in the CinePlex Basic Behavior module and also saved in a Plexon data file for further offline analysis. Based on the location of the rat, this system presented stimuli, collected behavioral data, and controlled delivery of reward. A OmniPlex Neural Data Acquisition System (Plexon, Inc) and SortClient (Plexon, Inc) recorded real-time neuronal activity and behaviorally relevant event timestamps for later analysis. The OminiPlex system was interfaced with the Med Associates system (DIG-716B SmartCTL, DIG-713A SuperPort TTL Input module and DIG-726 SuperPort TTL Output module).

### Behavioral training, surgery, and retraining

Prior to surgery, rats were habituated to the behavioral room and apparatus and then gradually shaped to the final parameters of the task. During shaping, the time in the ready position was gradually increased, the duration of the visual stimulus was gradually decreased, and the response window for selecting was gradually decreased to the final parameters. Shaping was accomplished in 9 to 11 weeks. In the final task, rats were required to stay in the ready position waiting for stimulus onset for a variable interval (1.2-1.6 seconds), at which point the stimulus appeared for 500 msec. Rats were required to make a selection by approaching the location of the stimulus within the 5 s response window. Accuracy was calculated as the number of correct trials divided by the total of correct and incorrect trials. A trial was defined as an omission trial when the rat did not make a selection during the response window. Prior to surgery, rats were trained on the final parameters to a criterion of 70-80% accuracy for at least five continuous days.

After reaching a behavioral criterion of performance, the rats were unilaterally implanted with a custom hyperdrive targeting the PUL, DPPC, and CPPC and counterbalanced across hemispheres (Figure 1C). Hyperdrives included fourteen stereotrode microdrives for recording neuronal activity and two fibers for optical suppression. In the same surgery, we injected a viral vector to transduce the unilateral PUL recording site with an inhibitory halorhodopsin (NpHR) driven by a synapsin 1 promoter packaged into an adeno-associated viral vector (AAV9). Histology verified that all stereotrodes were properly located and NpHR expression was observed in the PUL (Figure 1D).

Accuracy decreased after surgery and recovery but improved across sessions. Mean accuracy at the beginning of retraining was 55.0% (range 41.9%-67.1%), lower than pre-surgery levels, but still above chance level of 33.3%. Mean accuracy in the last sessions was 60.4% (range 44.7%-70.4%). The mean accuracy of the animal that was at 44.7% in the last session was 52.6%.

### Optical Suppression

PUL was transduced with the light sensitive inhibitory protein halorhodopsin. For optical suppression, we used multimode fiber (Paradigm optics; core diameter 240 microns, 0.5 NA). The fibers were pulled using a laser-based pipette puller (P-97; Sutter Instruments) and attached to a ferrule before insertion into the microdrive. All pulled fibers were tested before use and only fibers delivering light with at least 80% efficiency were used in the experiment. Light delivered by LED using a through a 4 channel optogenetic controller (PlexBright Optogenetic Stimulation System, Plexon, Inc) using Radiant software (Plexon, Inc.). The controller and connecting cable were tested before each session. On one-half of all trials, PUL neurons were stimulated optogenetically at 40 Hz with 15 ms pulse width. Stimulation began at trial onset and ended after the reward epoch. The LED light was shielded with electrical tape so as not to provide a distracting stimulus.

### Viral vectors

For viral transduction in the PUL, pAAV-hSyn-eNpHR 3.0-EYFP plasmid with an inhibitory halorhodopsin (NpHR)-EYFP fusion gene driven by a synapsin 1 promoter packaged into an adeno-associated viral vector (AAV9) at the University of Pennsylvania Vector Core was used. Plasmid maps are available at www.optogenetics.org. Viral titers were 2.04 e13 GC/ml.

### Behavioral training

Rats were put on food schedules to maintain body weight at 85-90% of free feeding weight. After handling for at least 7 days, rats were habituated to the behavioral room for 10 min/day for three days. Rats were then shaped and trained in the VSA task. In the shaping sessions, rats first were trained in a 30-minute session to approach a visual target stimulus for a food reward (a drop of flavored milk). In the initial shaping sessions, we adopted an errorless shaping procedure such that when the rat moved toward one of the three locations in one side of the maze, the visual stimulus at that location would illuminate and a tone would signal a correct choice. A new trial on the other side of the maze would be initiated after the rat entered reward area. After this initial shaping phase, rats were trained to stop in the ready position zone located in the middle of the bowtie shaped maze facing the stimulus presenting area. After a variable delay, a visual stimulus would illuminate in one of three randomly chosen locations. There was a short response window for rats to approach the location of the visual stimulus. Approach to the correct location was signaled by a brief tone and presentation of a drop of flavored milk as a food reward. If the rat approached an incorrect location, no reward was given, the trial was terminated, and a new trial would begin immediately. Rats were gradually trained in a series of steps culminating in the final parameters of the task. The duration for rats to stay in the ready position was gradually increased from 0.1 seconds to 1.6 seconds. Visual stimulus duration was gradually decreased from 20 seconds to 0.5 seconds. The response time window was decreased from 20 seconds to 5 seconds. In the final task, rats were required to stay in the ready position until stimulus onset for a variable interval (1.2-1.6 seconds) (Figure 1A).

The behavioral performance criterion for the end of training was 70-80% accuracy on the VSA task (chance is 33.33%). Behavior was well controlled on correct trials. Rats in this study required 2-3 months of training to reach the behavioral criterion on the final stage of the task. A surgery for implanting the hyperdrive was done after a rat reached behavioral criterion for 5 to 7 consecutive days. The response latencies were recorded for correct and incorrect trials and were calculated as follows: timestamp of selection – timestamp of start of stimulus onset. Latency to collect reward was calculated as follows: timestamp to enter the reward port location– timestamp of selection. We used a t-test to compare the response and reward latencies between correct and incorrect trials.

For recording sessions after implanting surgery, optical suppressions were added randomly to trials for whole session. The duration of optical suppression started when the rat entered ready position initiating a new trial and ended when the rat entered the reward zone terminating the trial. So, for one recording session, there were both trials with and without optical suppression (Figure 1B).

### Surgery

Animals were premedicated with diazepam (2-5 mg/kg; i.p.), glycopyrrolate (0.05 mg/kg; s.c.), carprofen (5 mg/kg; s.c.), and butorphanol tartrate (0.5 mg/kg; s.c.) to counteract respiratory effects of anesthesia, to control pain, and to decrease risk of seizures. They were brought to a surgical level of anesthesia with isofluorane (1.0 – 2.5%). Using a stereotaxic apparatus (Kopf, Tujunga, CA), rats were unilaterally implanted with a custom hyperdrive into the PUL (AP –3.9 mm, ML ± 2.0 mm, DV–4.0mm), DPPC (AP – 4.2 mm, ML ± 3.0mm, DV −0.1mm), and CPPC (AP – 5.5 mm, ML ± 4.5 mm, DV −2.0 mm). The hyperdrive had fourteen microdrives for recording neuronal activity and two fibers for optical suppression. Each microdrive consisted of a drivable screw with guide tubing containing on stereotrode. Stereotrodes were made of two 12 μm twisted, formvar-insulated nichrome wires (A-M systems, Sequim, WA). A full turn of the screw advanced the stereotrode by 350 μm. Two silver ground wires were wrapped around anchor screws in the skull. The hyperdrive was secured to the skull by the ground screws, small anchor screws, grip cement (Dentsply Caulk, Milford, DE), and dental cement (Coltene/Whaledent Inc., Cuyahoga Falls, OH).

### Histology

After the last recording session, the rats were deeply anesthetized with an overdose of Beuthanasia-D (100 mg/kg, i.p.) and the final recording site was marked with an electrolytic lesion. The rats were then perfused with normal saline, followed by 4% formalin. The brains were post-fixed for 24 hours in 4% formalin and then transferred to a 30% sucrose solution until the time for sectioning. The brains were sectioned at 40 μm. One set stained with Nissl material using thionin for examining electrode implanting sites. Another set stained for DAPI (VECTASHIELD, Vector Laboratories) to see the NpHR expression in the PUL.

### Electrophysiology

Neuronal activity recorded from stereotrodes, was amplified with a gain of 20 at the head stage (HST/32V-G20, Plexon, Inc., Dallas, TX). Signals were passed through a high-gain amplifier (total = 10000X, Omniplex system, Plexon, Inc., Dallas, TX). Single-unit activity was filtered between 0.8 - 6 Hz. The signal was then digitized at 40 kHz for single-unit activity. These signals were extracted through real-time thresholding (Sort Client, Plexon, Inc). The final waveforms were stored with timestamps of relevant events and position information for later analysis. Cells included in analyses had good quality waveforms (mean ± standard deviation) from both electrodes and were well clustered. Figure 1E shows a representative example of waveforms identified as belonging to an isolated cell from each region along with the corresponding autocorrelogram.

### Single neuron activity analysis

Spikes associated with putative individual cells were isolated offline based on waveform characteristics and using a variety of partially automated and manual techniques (Offline Sorter, Plexon, Inc.). The result was a dataset for each cell containing timestamps corresponding to spike times and behaviorally relevant event markers. These datasets were further analyzed using Neuroexplorer (NEX, Nex Technologies, Madison, AL), SPSS (IBM Corporation, Somers, NY) and Matlab (Mathworks, Natick, MA).

#### Analysis of variance (ANOVA)

In our first set of analysis, each cell was analyzed for behavioral correlates using a two-way ANOVA. We analyzed five epochs from each completed trial: pre-stimulus and post-stimulus epochs were the 500 msec periods immediately before and after stimulus onset; pre-selection and post-selection epochs were the 500 msec periods immediately before and after the rat selected a target by approaching the location; and the reward-approach epoch was the first 500 msec after the animal entered the food reward area. The selection time event was the moment the rat entered a zone just in front of the location in which the target stimulus had appeared. Entry of the ready position of the other side of bowtie maze triggered the next trial. Rats invariably approached and checked the food port even after incorrect trials.

FR was the dependent variable for the ANOVAs. For each neuron, we first computed the mean FR (spikes/sec) for each epoch on each trial. In the first set of analyses, the between-trial variable was outcome (correct response vs. incorrect response), and the two within-trial variables were stimulus onset (pre-stimulus vs. post-stimulus) and selection time (pre-selection vs. post-selection). Outcome was analyzed for the reward-approach epoch.

In a second set of analyses, we examined neural correlates associated with the location of the target stimulus using one-way ANOVA. Based on the location of stimulus presentation, we pooled trials in which the target stimulus was at the same side of the maze (east vs. west) for analyzing allocentric location correlates. We then pooled trials in which the target stimulus was at the same egocentric location (left, right, and center) for analyzing egocentric location correlates. An ANOVA was computed on each cell from all target regions. Both allocentric location and egocentric location were analyzed separately for the post-stimulus, pre-selection, and post-selection epochs.

#### Sliding window analysis

To further understand changes of neuronal activity over time, a sliding window analysis was employed. We focused on the neuronal activity during the stimulus presentation period. For each neuron, we took a 100 ms window of time, beginning at stimulus onset and compared the mean FR of this 100 ms bin to the neuron’s mean FR during pre-stimulus epoch by using paired t-test. We then advanced the window by 20 ms and analyzed the next 100 ms window of time. We continued that process until the end of the stimulus presentation. Significance of a cell was determined by at least two continuous bins showing statistically significant differences (p < 0.05).

We then calculated the difference between the two conditions in the latency for cells to become selective: LED-OFF time to respond minus LED-ON time to respond. Thus, negative values indicate the cell became selective earlier, zero means the cell became selective in the same window in both conditions, and positive values indicate the cell became selective later. We then counted the number of cells in 20 ms windows from −380 to +380. We then conducted pairwise comparisons of the percentage distributions using the Kolmogorov-Smirnov test, and also calculated the percentages of earlier, same, and later selectivity change and subjected those distributions to pairwise Chi-Square analyses, Bonferroni corrected for multiple comparisons.

#### Multiple linear regression

We used multiple linear regression to examine the effects of optical suppression on neural activity at each of five 500-ms epochs (pre-stimulus, post-stimulus, pre-selection, post-selection, and reward), we first created full and reduced regression models (Equations #1 and #2, respectively) :

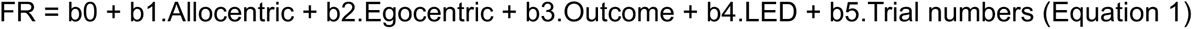

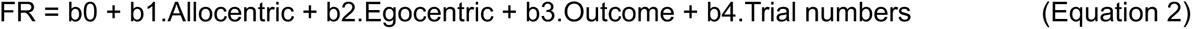

The full model included five variables to predict the mean FR of each neuron in each epoch. The allocentric predictor indicated the location in allocentric space where the rat expected the target to be illuminated regardless of the rat’s location or viewpoint and consisted of 2 levels (−1 for west trials, 1 for east trials). The egocentric predictor indicated the location of the illuminated target in egocentric space as the rat faced the expected allocentric location (i.e., left, center, or right, regardless of east or west location), and consisted of three levels (−1 for the location contralateral to the recorded hemisphere, 0 for the middle location, and 1 for the location ipsilateral to the recorded hemisphere). The outcome predictor had two levels (0 for incorrect responses, 1 for correct responses), and the LED predictor also had two levels (0 for LED-OFF trials, 1 for LED-ON trials). The trial number predictor was included as a nuisance parameter that would capture gradual changes in the neuron’s FR across the course of the recording session. Significance was evaluated at p < 0.05 unless otherwise indicated. Mean FR and predictors were standardized to allow comparison of regression coefficients. In the reduced model, we separated all response trials into two groups, LED-ON trials and LED-OFF trials, and used the remaining variables to predict the mean FR within each trial group at each epoch. In addition, we calculated the coefficient of partial determination (CPD) for each group in sliding time windows (100-ms window size and 20-ms steps), within 1000-ms before/after stimulus and selection onsets. CPD is the amount of variance in the neuron’s FR that can be explained by one predictor over and above the variance explained by other predictors included in the model ^40^. The CPD for predictor i is defined as

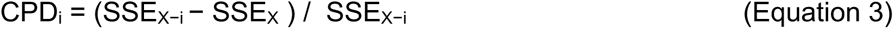

where SSE_X−i_ is the sum of squared errors in a regression model that includes all of the relevant predictor variables except predictor i, and SSE_X_ is the sum of squared errors in a regression model that includes all of the relevant predictor variables. To examine the time course of the contribution of each predictor, we performed a sliding regression analysis to calculate the CPD at each time point for each neuron. The sliding regression analysis requires a correction for multiple comparisons, because it involves performing a statistical test for multiple overlapping time points. We calculated this correction by computing a false alarm rate using shuffled neural data where significant parameters can only reflect noise. We preserved the experimental conditions (the design matrix) but shuffled the trial order of mean FR 1000 times randomly to create a null distribution for each neuron. CPD_i_ for a given time window was defined as significant if the actual CPD was larger than the 95^th^ percentile of null distribution. Once the significance of each CPD window was decided, a response latency curve was created for a neural population by sorting the neurons according to the earliest time with at least three consecutive significant windows. The non-significant windows were color-labeled in white for data visualization.

#### Decoding analysis

Linear determinant analysis (LDA) was used to decode six cue locations from neural activity using Matlab function *classify* (MathWorks, Inc). Considering the single implant side of the optical fiber on each subject, cue locations were labeled as ipsilateral, middle, and contralateral to the optical fiber during west or east trials. This label arrangement allowed us to take implant location into account in the analysis of allocentric and egocentric information while subjects navigated the floor maze continuously. Because numbers of recorded units and LED-ON/OFF trials were not exactly the same across recording sessions and subjects, we instituted criteria for data to be included in the decoding analysis. To have recorded units with full firing patterns in each location, we first included in the analysis only units from recording sessions with at least 3 correct trials in LED-ON and/or LED-OFF trials in each cue location. Units in any session that failed to reach this criterion were not included in the pseudo-populations. In addition, we excluded the recorded units from rat #4 in the decoding analyses because the units with higher firing rates (12 units from PUL, 9 units from DPPC, and 9 units from CPPC) were found specifically at the west-contralateral location during LED-ON trials, leading to the obvious distortions when the method of dimensionality reduction was applied. Overall, these preprocessing steps allowed us to apply cross condition decoding corresponding to the three types of train-test conditions. Accordingly, we included 240, 79, and 109 units from PUL, DPPC, and CPPC in train OFF & test OFF condition. Also, we included 116, 41, and 62 units from PUL, DPPC, and CPPC in train OFF & test ON condition. Lastly, we included 145, 60, and 75 units from PUL, DPPC, and CPPC in train ON & test ON condition (Table S4). To ensure the number of trials in the feature matrix were equal across units, we found the minimum trial number (*n*) for each of 6 cue locations in the pseudo-population. Next, we randomly selected *n* trials from each cue location for inclusion in the analysis. Once the feature matrix (e.g. 18 trials-by-240 units) and the vector of cue locations were decided, we ran principal component analysis (PCA) and kept the first 5 PCs, and these were submitted to LDA with leave-one-trial-out cross validation^34^ ^41^. We repeated the randomized trial sampling procedure 1000 times to ensure that all trials are included in the decoding analyses and computed the mean posterior probability with standard error mean (SEM) for each cue location, as well as the mean decoding performance for comparing decoding with or without optical suppression.

#### Distributions of projected firing patterns to PCA subspace

To further determine whether the firing patterns of recorded units were influenced by optical suppression of PUL, we assessed firing patterns of train OFF & test ON condition used in the decoding analyses above. For each feature matrix established by the randomized trial sampling procedure, we projected the trial data points as the firing patterns from training (LED-OFF trials) and testing (LED-ON trials) groups onto the first 5 PCs (Figure S4). We calculated three measurements to describe how those two groups are distributed. First, we found the centroid for each group of projected trials based on Matlab function *kmeans* with the cosine distances and calculated the vector length from the origin to the centroid (Figure S4A and S4C). Second, we found the dispersion of data points around the centroid by calculating the cluster variance, with higher variances indicating that the data points were more distributed away from the centroid (Figure S4B). Lastly, we calculated the vector angles between the centroids of training and testing groups (Figure S4D). If optical suppression influenced firing patterns, then we expected that the two groups of data points (trials) would have different centroids in PC subspace, and therefore large vector angles. One or paired-sample t-test were computed to compare the difference of measurements between training and testing groups (p < 0.05).

## SUPPLEMENTAL MATERIALS

### Supplementary Figures

**Supplemental Figure S1. for Figure 3, Panels D-F.**
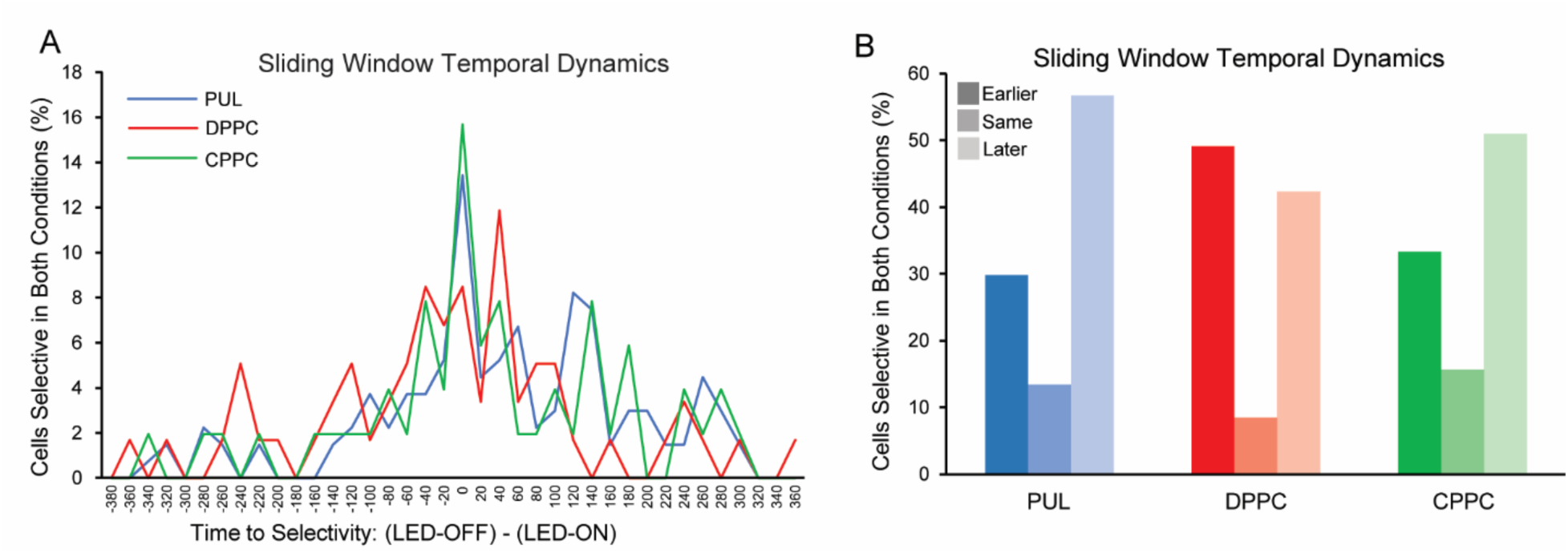
Sliding window analysis of temporal dynamics of selectivity onset. For cells that were selective in both LED-OFF and LED-ON, the difference in the latency to onset selectivity in the two conditions was calculated as LED-OFF minus LED-On. Cells either became selective later (negative values), in the same 20 ms window (0 point), or earlier (positive values). The x-axis indicates the start point of each 20 ms window. Thus, the first window is from −380 to −360 and the last window is from 360 to 380. (A) Shown are percentage distributions of cells according to the difference calculation. The distributions for DPPC and CPPC were significantly different. (B) Percentages of cells that showed negative, 0, or positive values. For PUL and CPPC, more cells showed later selectivity than earlier selectivity. The opposite pattern was evident for DPPC. Distributions for both PUL and CPPC were significantly different from the DPPC distribution, and they were not different from each other.

**Supplemental Figure S2. for Figure 6.**
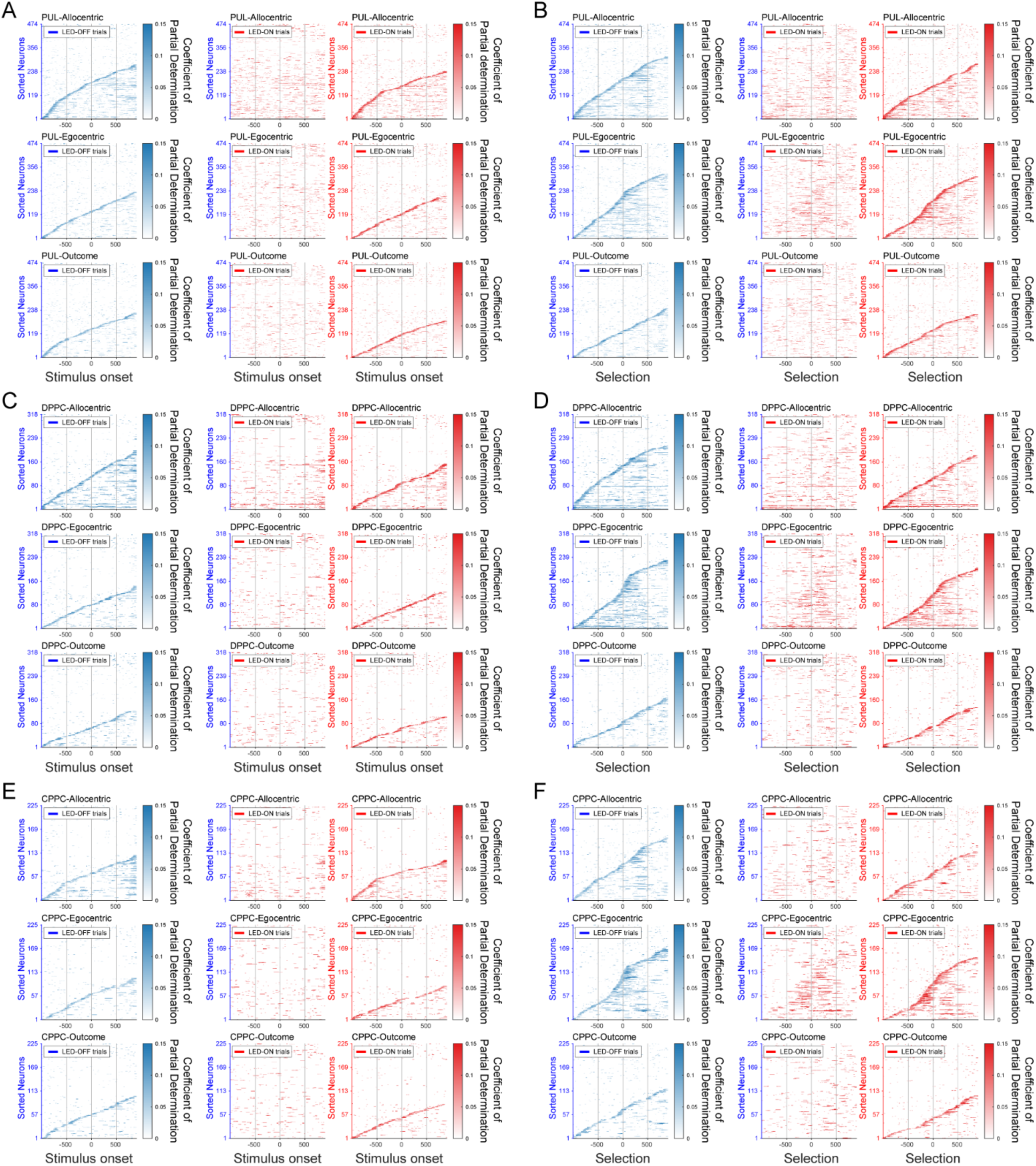
Time courses of coefficient of partial determination (CPD). Time courses of CPD used to produce latency curves for three predictors in LED-OFF (blue) and LED-ON (red) trials at two epochs of stimulus onset and selection onset in PUL (A and B), DPPC (C and D), and CPPC (E and F).

**Supplemental Figure S3. for Figure 7 and Figure 8.**
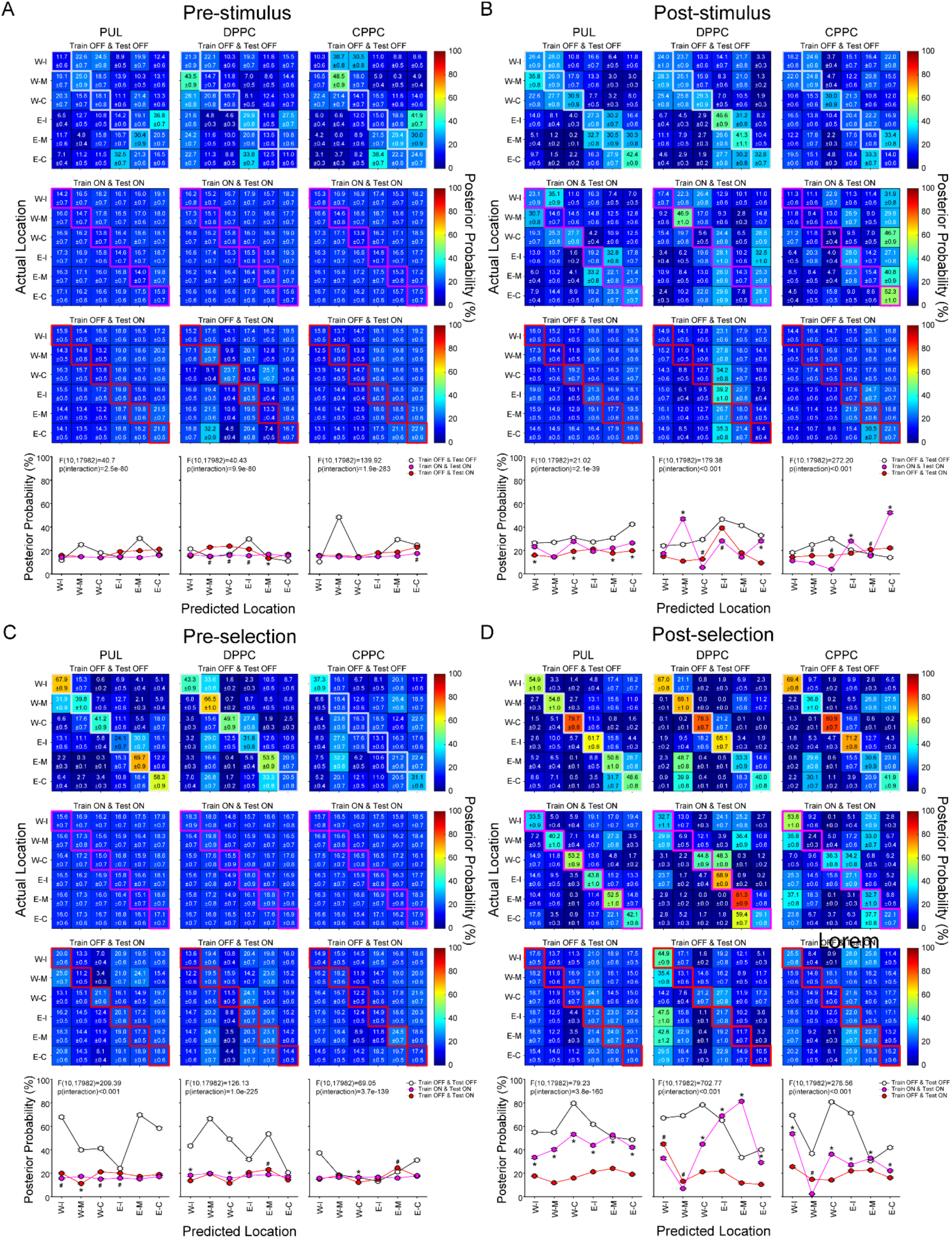
Summaries of decoding posterior probabilities in additional epochs. Heat plots show results from three train & test conditions in PUL, DPPC and CPPC during the pre-stimulus epoch (A), post-stimulus epoch (B), pre-selection epoch (C), and post-selection epoch (D), summarized in the bottom row as in Figure 7. Significant interactions of two-way ANOVA were listed in each subplots, and p-values for post-hoc comparisons were corrected by false discovery rate (FDR). * = posterior probabilities in Train ON & Test ON are significantly larger than those in Train OFF & Test ON (p < 0.001). ^#^ = posterior probabilities in Train OFF & Test ON are significantly larger than those in Train ON & Test ON.

**Supplemental Figure S4.**
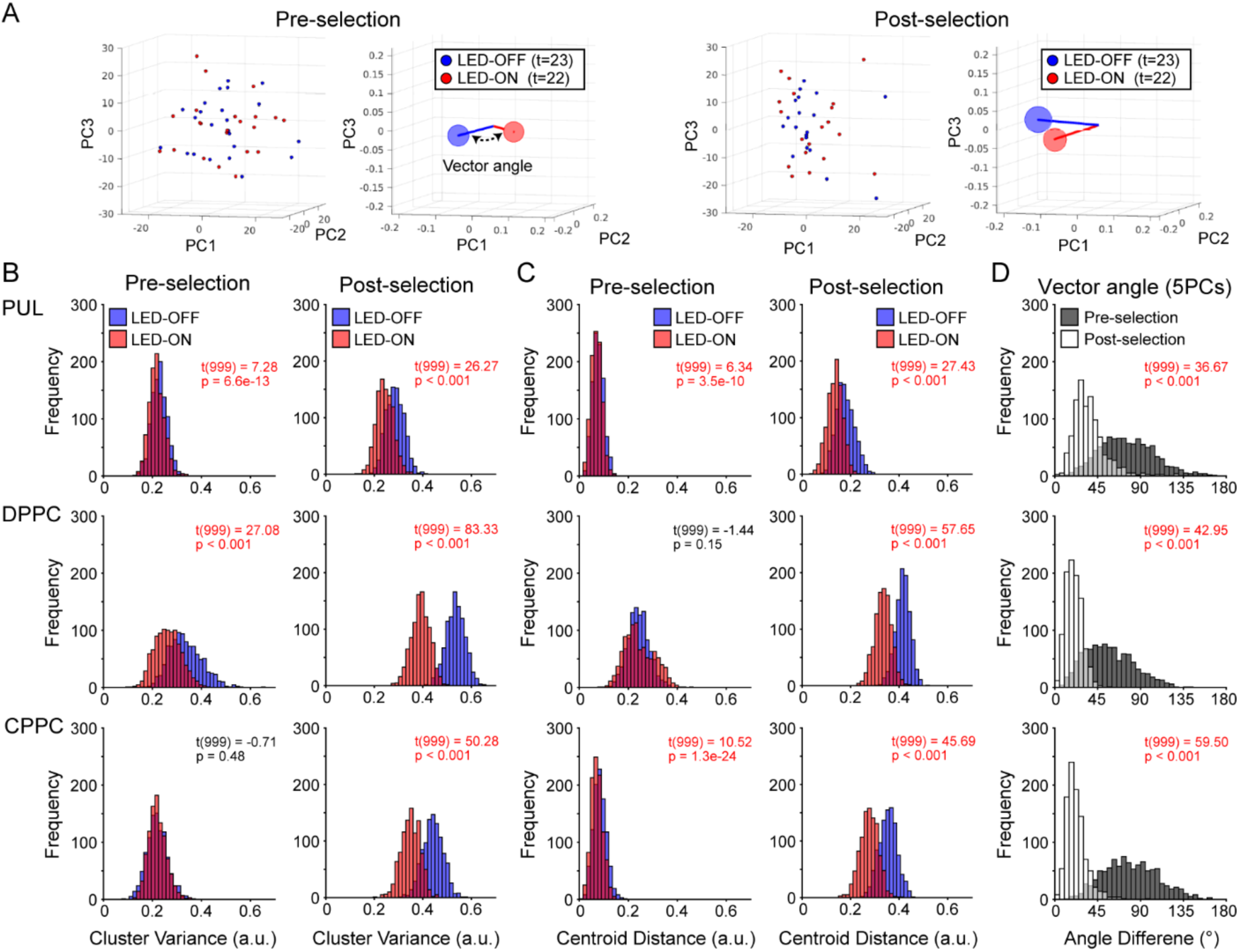
LED-OFF and LED-ON trials occupy different subspaces. (A) A single sampling dataset from the PUL pseudopopulation shown on the first 3 PCs for visualization (actual results calculated from the first 5 PCs). Two sample epochs, pre- and post-selection, are shown. The left panel at each epoch shows individual pseudotrials in LED-OFF (blue, 23 trials) and LED-ON (red, 22 trials) groups, and the right panels show the centroids and vectors to the centroids (calculated by k-means with cosine distance) for both groups. Dispersion of the pseudotrials was estimated by centroid variances and illustrated as the relative size of circular markers. The vector distance from the origin to the center of each centroid also characterized the reliability of the PC loadings across trials. Lastly, vector angles between two centroids quantified the separation of the two conditions. (B) Distributions of the cluster variances. LED-OFF (blue) and LED-ON (red) trials shown in each panel were compared with one sample t-tests. Significant results were color-labeled in red (p < 0.001). LED-OFF trials tended to have higher cluster variance, especially in the post-selection epoch, suggesting more trial-wise variance in LED-OFF trials. Results from PUL, DPPC, and CPPC areas in rows. (C) The distributions of the vector lengths between two centroid in two epochs. There were again larger vector lengths during the LED-OFF condition. Legend captions described as (B). (D) The distributions of the angles between two centroid vectors in two epochs. Angles were generally between 45 and 90 degrees in pre-selection (gray) epoch, but less than 45 degrees in post-selection (white) epoch, consistent with a change in firing patterns across LED conditions in (B) and (C). Further, significant differences across epochs were found in each area before/after target selections, indicating that the information from PUL to DPPC and CPPC may alter the differences of ensemble firing after target selections with optical perturbation. Statistical tests described as (B).

### Supplementary Tables

**Table S1.** Cells selective for peri-stimulus and peri-selection epochs in LED OFF and LED-ON conditions. Supplementary Table for Figure 3, A-C.

| Epoch | Correlate | PUL (n = 474)<br>Number (%) |  | DPPC (n = 318)<br>Number (%) |  | CPPC (n= 225)<br>Number (%) |  |
| --- | --- | --- | --- | --- | --- | --- | --- |
|  |  | LED-OFF | LED-ON | LED-OFF | LED-ON | LED-OFF | LED-ON |
| Peri-Stimulus | Stimulus Only | 78 (16.5) | 64 (13.5) | 34 (10.7) | 21 (6.6) | 24 (10.7) | 21 (9.3) |
|  | Stimulus and Outcome | 35 (7.4) | 34 (7.2) | 16 (5.0) | 12 (3.8) | 32 (14.2) | 13 (5.8) |
|  | Outcome Only | 17(3.6) | 19 (4.0) | 18 (5.7) | 12 (3.8) | 9 (4.0) | 13 (5.8) |
|  | Total | 130 (27.4) | 117 (24.7) | 68 (21.4) | 45 (14.2) | 65 (28.9) | 47 (20.9) |
| Peri-Selection | Selection Only | 160 (33.8) | 106 (22.4) | 122 (38.4) | 91 (28.6) | 98 (43.6) | 63 (28.0) |
|  | Selection and Outcome | 30 (6.3) | 46 (9.7) | 24 (7.5) | 24 (7.5) | 20 (8.9) | 25 (11.1) |
|  | Outcome Only | 21 (4.4) | 15 (3.2) | 10 (3.1) | 7 (2.2) | 6 (2.7) | 7 (3.1) |
|  | Total | 211 (44.5) | 167 (35.2) | 156 (49.1) | 122 (38.4) | 124 (55.1) | 95 (42.2) |
| Reward | Outcome | 59 (12.4) | 45 (9.5) | 38 (11.9) | 20 (6.3) | 22 (9.8) | 13 (5.8) |
|  | Nonselective | 415 (87.6) | 429 (90.5) | 280 (88.1) | 298 (93.7) | 203 (90.2) | 212 (94.2) |

Note that some cells are selective for multiple correlates such that the totals are less than the sum of correlates.

**Table S2.**
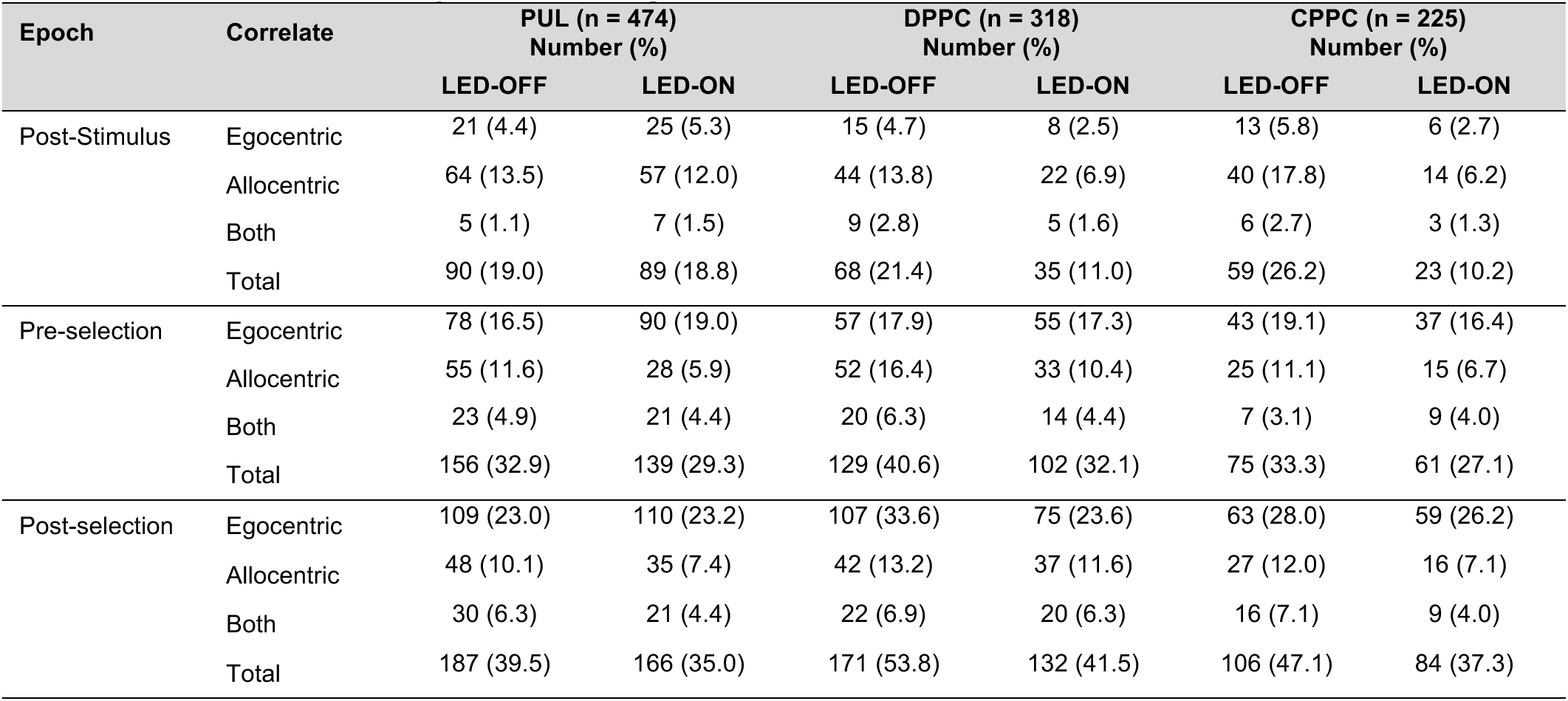
Cells selective for allocentric and egocentric locations in LED OFF and LED-ON conditions. Supplementary Table for Figure 3, G-I.

**Table S3.** Changes in Selectivity between LED-OFF and LED-ON trials. Supplementary table for Figure 4.

| LED Impact on correlates of stimulus onset and outcome |  |  |  |  |  |
| --- | --- | --- | --- | --- | --- |
| Peri-Stimulus | LED-OFF only | LED-ON only | Off-On Altered Selectivity | Off-On Same Selectivity | Total Reorganized |
| PUL (n = 188, 40%) | 71 (37.8) | 58 (30.9) | 26 (13.8) | 33 (17.6) | 155 (82.4) |
| DPPC (n = 102, 32%) | 57 (55.9) | 34 (33.3) | 3 (2.9) | 8 (7.8) | 94 (92.2) |
| CPPC (n = 90, 40%) | 43 (47.8) | 25 (27.8) | 10 (11.1) | 12 (13.3) | 78 (86.7) |
| Sliding Window | LED-OFF only | LED-ON only | Off-On Altered Latency to Selectivity | Off-On Same Latency to Selectivity | Total Reorganized |
| PUL (n = 310, 65%) | 112 (36.1) | 85 (27.4) | 18 (5.8) | 95 (30.7) | 215 (69.4) |
| DPPC (n = 170, 53%) | 59 (34.7) | 52 (30.6) | 5 (2.9) | 54 (31.8) | 116 (68.2) |
| CPPC (n = 144, 64%) | 60 (41.7) | 32 (22.2) | 8 (5.6) | 44 (30.6) | 100 (69.4) |
| LED Impact on correlates of decision making and outcome |  |  |  |  |  |
| Peri-Selection | LED-OFF only | LED-ON only | Off-On Altered Selectivity | Off-On Same Selectivity | Total Reorganized |
| PUL (n = 270, 57%) | 103 (38.1) | 59 (21.9) | 42 (15.6) | 66 (24.4) | 204 (75.6) |
| DPPC (n = 193, 61%) | 71 (36.8) | 37 (19.2) | 30 (15.5) | 55 (28.5) | 138 (71.5) |
| CPPC (n = 151, 67%) | 57 (37.7) | 28 (18.5) | 20 (13.3) | 46 (30.5) | 105 (69.5) |
| LED Impact on coding for egocentric and allocentric reference frames |  |  |  |  |  |
| Epoch | LED-OFF only | LED-ON only | Off-On Altered Selectivity | Off-On Same Selectivity | Total Reorganized |
| **Post-stimulus** |  |  |  |  |  |
| PUL (n = 152, 32%) | 62 (40.8) | 62 (40.8) | 10 (6.6) | 18 (11.8) | 134 (88.2) |
| DPPC (n = 84, 26%) | 49 (58.3) | 16 (19.0) | 9 (10.7) | 10 (11.9) | 74 (88.1) |
| CPPC (n = 76, 34%) | 53 (69.7) | 17 (22.4) | 4 (5.3) | 2 (2.6) | 74 (97.4) |
| **Pre-selection** |  |  |  |  |  |
| PUL (n = 197, 42%) | 76 (38.6) | 58 (29.4) | 27 (13.7) | 36 (18.3) | 161 (81.7) |
| DPPC (n = 160, 50%) | 58 (36.3) | 31 (19.4) | 44 (27.5) | 27 (16.9) | 133 (83.1) |
| CPPC (n = 98, 44%) | 37 (37.8) | 23 (23.5) | 13 (13.3) | 25 (25.5) | 73 (74.5) |
| **Post-selection** |  |  |  |  |  |
| PUL (n = 251, 53%) | 86 (34.3) | 65 (25.9) | 48 (19.1) | 52 (20.7) | 199 (79.3) |
| DPPC (n = 198, 62%) | 66 (33.3) | 27 (13.6) | 63 (31.8) | 42 (21.2) | 156 (78.8) |
| CPPC (n = 125, 56%) | 41 (32.8) | 19 (15.2) | 27 (21.6) | 38 (30.4) | 87 (69.6) |

Shown in left column are regions and numbers of cells showing selectivity in at least one condition. Percent is the percent of total isolated cells (474 in the PUL, 318 in the DPPC and 225 in the CPPC ). Other columns show Number (% selective). Total reorganized are total cells that responded to LED-ON by losing, gaining, or changing selectivity.

**Table S4.**
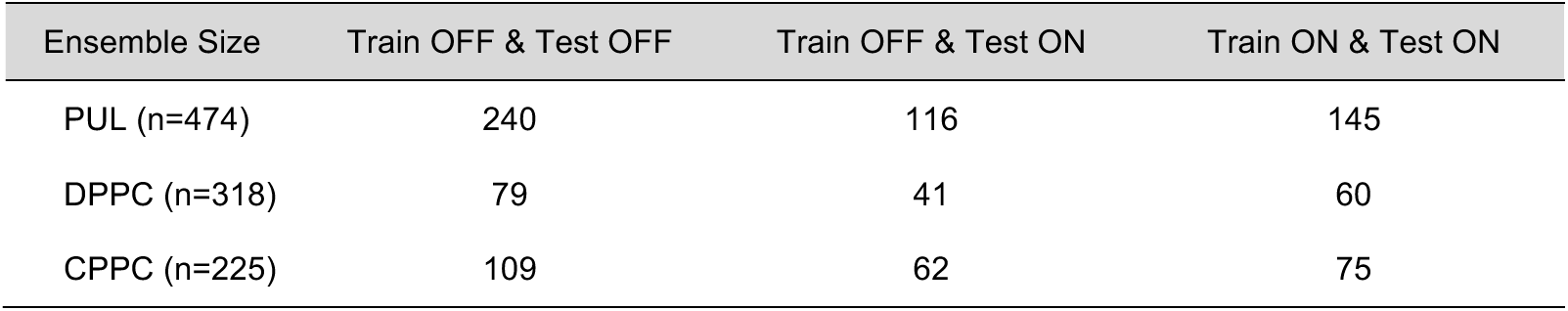
Ensemble size for cross condition decoding corresponding to three types of train-test conditions. Supplementary table for Figure 7 and Figure 8.

## References

1. Scholl, L.R., Foik, A.T., and Lyon, D.C. (2021). Projections between visual cortex and pulvinar in the rat. Journal of Comparative Neurology 529, 129–140. 10.1002/cne.24937.

2. Takahashi, T. (1985). The organization of the lateral thalamus of the hooded rat. Journal of Comparative Neurology 231, 281–309. 10.1002/cne.902310302.

3. Guillery, R.W. (1995). Anatomical evidence concerning the role of the thalamus in corticocortical communication: a brief review. Journal of anatomy 187 ( *Pt 3*), 583–592.

4. Yang, F.-C., Dokovna, L.B., and Burwell, R.D. (2022). Functional differentiation of dorsal and ventral posterior parietal cortex of the rat: Implications for controlled and stimulus-driven attention. Cerebral Cortex 32, 1787–1803.

5. Kaas, J.H., and Lyon, D.C. (2007). Pulvinar contributions to the dorsal and ventral streams of visual processing in primates. Brain Research Reviews 55, 285–296. 10.1016/j.brainresrev.2007.02.008.

6. Chandler, H.C., King, V., Corwin, J.V., and Reep, R.L. (1992). Thalamocortical connections of rat posterior parietal cortex. Neuroscience Letters 143, 237–242. 10.1016/0304-3940(92)90273-A.

7. Reep, R.L., Chandler, H.C., King, V., and Corwin, J.V. (1994). Rat posterior parietal cortex: topography of corticocortical and thalamic connections. Experimental Brain Research 100, 67–84.

8. Fiebelkorn, I.C., and Kastner, S. (2019). The puzzling pulvinar. Neuron 101, 201–203. 10.1016/j.neuron.2018.12.032.

9. Halassa, M.M., and Sherman, S.M. (2019). Thalamocortical circuit motifs: A general framework. Neuron 103, 762–770. 10.1016/j.neuron.2019.06.005.

10. Shine, J.M., Lewis, L.D., Garrett, D.D., and Hwang, K. (2023). The impact of the human thalamus on brain-wide information processing. Nature Reviews Neuroscience. 10.1038/s41583-023-00701-0.

11. Halassa, M.M., and Kastner, S. (2017). Thalamic functions in distributed cognitive control. Nature Neuroscience 20, 1669–1679. 10.1038/s41593-017-0020-1.

12. Yang, F.-C., and Burwell, R.D. (2020). Neuronal activity in the rat pulvinar correlates with multiple higher-order cognitive functions. Vision 4, 15.

13. Reep, R.L., and Corwin, J.V. (2009). Posterior parietal cortex as part of a neural network for directed attention in rats. Neurobiology of Learning and Memory 91, 104–113. 10.1016/j.nlm.2008.08.010.

14. Karnath, H.O., Himmelbach, M., and Rorden, C. (2002). The subcortical anatomy of human spatial neglect: putamen, caudate nucleus and pulvinar. Brain 125, 350–360. 10.1093/brain/awf032.

15. Snow, J.C., Allen, H.A., Rafal, R.D., and Humphreys, G.W. (2009). Impaired attentional selection following lesions to human pulvinar: Evidence for homology between human and monkey. Proceedings of the National Academy of Sciences 106, 4054–4059.

16. Behrmann, M., Geng, J.J., and Shomstein, S. (2004). Parietal cortex and attention. Current Opinion in Neurobiology 14, 212–217. 10.1016/j.conb.2004.03.012.

17. Kravitz, D.J., Saleem, K.S., Baker, C.I., and Mishkin, M. (2011). A new neural framework for visuospatial processing. Nature Reviews Neuroscience 12, 217–230.

18. Broussard, J., Sarter, M., and Givens, B. (2006). Neuronal correlates of signal detection in the posterior parietal cortex of rats performing a sustained attention task. Neuroscience 143, 407–417. 10.1016/j.neuroscience.2006.08.030.

19. Nitz, D.A. (2012). Spaces within spaces: rat parietal cortex neurons register position across three reference frames. Nature Neuroscience 15, 1365–1367.

20. Yang, F.-C., Jacobson, T.K., and Burwell, R.D. (2017). Single neuron activity and theta modulation in the posterior parietal cortex in a visuospatial attention task. Hippocampus 27, 263–273. 10.1002/hipo.22691.

21. Nitz, D.A. (2006). Tracking route progression in the posterior parietal cortex. Neuron 49, 747–756. 10.1016/j.neuron.2006.01.037.

22. Wilber, A.A., Clark, B.J., Demecha, A.J., Mesina, L., Vos, J.M., and McNaughton, B.L. (2015). Cortical connectivity maps reveal anatomically distinct areas in the parietal cortex of the rat. Frontiers In Neural Circuits 8. 10.3389/fncir.2014.00146.

23. Olsen, G.M., and Witter, M.P. (2016). Posterior parietal cortex of the rat: Architectural delineation and thalamic differentiation. Journal of Comparative Neurology 524, 3774–3809. 10.1002/cne.24032.

24. Corbetta, M., and Shulman, G.L. (2002). Control of goal-directed and stimulus-driven attention in the brain. Nature Reviews Neuroscience 3, 201–215.

25. Shomstein, S. (2012). Cognitive functions of the posterior parietal cortex: top-down and bottom-up attentional control. Frontiers in Integrative Neuroscience 6, 38.

26. Carli, M., Robbins, T.W., Evenden, J.L., and Everitt, B.J. (1983). Effects of lesions to ascending noradrenergic neurones on performance of a 5-choice serial reaction task in rats; implications for theories of dorsal noradrenergic bundle function based on selective attention and arousal. Behavioural Brain Research 9, 361–380. 10.1016/0166-4328(83)90138-9.

27. Furtak, S.C., Cho, C.E., Kerr, K.M., Barredo, J.L., Alleyne, J.E., Patterson, Y.R., and Burwell, R.D. (2009). The Floor Projection Maze: A novel behavioral apparatus for presenting visual stimuli to rats. Journal of Neuroscience Methods 181, 82–88. 10.1016/j.jneumeth.2009.04.023.

28. Jacobson, T.K., Ho, J.W., Kent, B.W., Yang, F.-C., and Burwell, R.D. (2014). Automated visual cognitive tasks for recording neural activity using a floor projection maze. JoVE, e51316. doi:10.3791/51316.

29. Parthasarathy, A., Tang, C., Herikstad, R., Cheong, L.F., Yen, S.-C., and Libedinsky, C. (2019). Time-invariant working memory representations in the presence of code-morphing in the lateral prefrontal cortex. Nature Communications 10, 4995. 10.1038/s41467-019-12841-y.

30. Ciaramelli, E., and Moscovitch, M. (2020). The space for memory in posterior parietal cortex: Re-analyses of bottom-up attention data. Neuropsychologia 146, 107551. 10.1016/j.neuropsychologia.2020.107551.

31. Johnston, W.J., Palmer, S.E., and Freedman, D.J. (2020). Nonlinear mixed selectivity supports reliable neural computation. PLOS Computational Biology 16, e1007544. 10.1371/journal.pcbi.1007544.

32. Libedinsky, C. (2023). Comparing representations and computations in single neurons versus neural networks. Trends in Cognitive Sciences 27, 517–527. 10.1016/j.tics.2023.03.002.

33. Danziger, S., Ward, R., Owen, V., and Rafal, R. (1999). The effects of unilateral pulvinar damage in humans on reflexive orienting and filtering of irrelevant information. Behavioural Neurology 13, 95–104.

34. Chiang, F.-K., Wallis, J.D., and Rich, E.L. (2022). Cognitive strategies shift information from single neurons to populations in prefrontal cortex. Neuron 110, 709–721.e704. 10.1016/j.neuron.2021.11.021.

35. Agster, K.L., and Burwell, R.D. (2009). Cortical efferents of the perirhinal, postrhinal, and entorhinal cortices of the rat. Hippocampus 19, 1159–1186. 10.1002/hipo.20578.

36. Furtak, Sharon C., Ahmed, Omar J., and Burwell, Rebecca D. (2012). Single neuron activity and theta modulation in postrhinal cortex during visual object discrimination. Neuron 76, 976–988. 10.1016/j.neuron.2012.10.039.

37. Saalmann, Y.B., Pinsk, M.A., Wang, L., Li, X., and Kastner, S. (2012). The pulvinar regulates information transmission between cortical areas based on attention demands. Science 337, 753–756. 10.1126/science.1223082.

38. Zhou, H., Schafer, Robert J., and Desimone, R. (2016). Pulvinar-cortex interactions in vision and attention. Neuron 89, 209–220. 10.1016/j.neuron.2015.11.034.

39. Fiebelkorn, I.C., Pinsk, M.A., and Kastner, S. (2019). The mediodorsal pulvinar coordinates the macaque fronto-parietal network during rhythmic spatial attention. Nature Communications 10, 215. 10.1038/s41467-018-08151-4.

40. Chiang, F.-K., and Wallis, J.D. (2018). Neuronal encoding in prefrontal cortex during hierarchical reinforcement learning. Journal of Cognitive Neuroscience 30, 1197–1208.

41. Chien, J.M., Wallis, J.D., and Rich, E.L. (2023). Abstraction of reward context facilitates relative reward coding in neural populations of the macaque anterior cingulate cortex. The Journal of Neuroscience 43, 5944. 10.1523/JNEUROSCI.0292-23.2023.

